# Oncogenic Ras-driven Dorsal/NF-κB signaling contributes to tumorigenesis in a *Drosophila* carcinoma model

**DOI:** 10.1101/2024.05.08.593126

**Authors:** Caroline Dillard, Jose Teles Reis, Ashish Jain, Roland Le Borgne, Heinrich Jasper, Tor Erik Rusten

## Abstract

Cancer-driving mutations synergize with inflammatory stress signaling pathways during carcinogenesis. *Drosophila melanogaster* tumour models are increasingly recognized as models to inform conserved molecular mechanisms of tumorigenesis with both local and systemic effects of cancer. Although initial discoveries of the Toll-NFκB signaling pathway in development and immunity was pioneered in *Drosophila*, limited information is available for its role in cancer progression. Using a well-studied cooperative Ras^V12^ -driven epithelial-derived tumour model, we here describe functions of Toll-NF-κB signaling in malignant *Ras^V12^, scrib^-^* tumors. The extracellular Toll pathway components ModSP and PGRP-SA and intracellular signaling Kinase, Pelle/IRAK, are rate-limiting for tumor growth. The Toll pathway NFκB protein Dorsal, as well as *cactus/I*κB show elevated expression in tumors with highest expression in invasive cell populations. Oncogenic Ras^V12^, and not loss of *scribble,* confers increased expression and heterogenous distribution of two Dorsal isoforms, DorsalA and DorsalB in different tumour cell populations. Mechanistic analyses demonstrates that Dorsal drives growth and malignancy by suppressing differentiation, counteracting apoptosis and promoting invasion of *Ras^V12^, scrib^-^* tumors genetically dependent on *twist* and *snail*.

## Introduction

Inflammation, defined as the body’s response to harmful external inputs, is a hallmark of cancer [1]. This response is mediated by four major inflammatory pathways: the MAPK, PI3K-AKT, JAK-STAT and NF-κB (canonical and non-canonical) signaling pathways which integrate intra- and extracellular alarms and trigger cellular responses [2]. NF-κB activation has been observed in many cancers [3, 4] and correlates with a poor prognosis in breast and non-small cell lung cancer [5, 6]. NF-κB functions appear cancer-relevant, as a significant body of *in vitro* studies reports its involvement in cellular processes such as invasion [7–9], proliferation [10, 11], stemness [12], survival [10–12] and chemoresistance [12–15]. Similarly, a limited number of human cancer xenograft studies demonstrated a positive effect of NF-κB on cancer proliferation [11], survival [11] and metastatic behavior [8, 9].

Despite several attempts, pharmaceutical targeting of NF-κB in cancer has proven ineffective due to toxicity resulting from its essential role in regulating cellular and innate immunity at the organismal level [16, 17]. Therefore, understanding both the upstream and downstream mechanisms governing NF-κB activation and functions in cancer, as well as deciphering whether such mechanisms are specific to cancer cells, is crucial. This understanding may facilitate the design of NF-κB -targeting cancer therapies that spare healthy cells. *In vivo* studies would be informative for achieving this goal.

*Drosophila melanogaster* is increasingly being used to study both cancer cell-autonomous mechanisms as well as the biology of complex local and systemic paraneoplastic effects instigated by tumor growth [18]. Among the *Drosophila melanogaster* cancer models, the *Ras^V12^, scrib^-/-^* carcinoma model, arising from the concomitant expression of the constitute activated oncogenic form of the KRAS/NRAS orthologue, Ras85D (here referred to as Ras^V12^), and the loss of cell polarity through *scribble* loss-of-function, has been most extensively studied and displays a remarkable array of hallmarks similar to human cancer [19] [20]. In *Ras^V12^, scrib^-/-^* tumors, generated in the larval eye-antennal disc epithelium (EAD), the evolutionarily conserved Ras-MAPK, PI3K-AKT and JAK-STAT inflammatory pathways all constitute major drivers [19, 20]. Surprisingly, at the organismal level, *Ras^V12^, scrib^-/-^*tumour-bearing flies also display both microenvironmental and systemic alterations that mimic human cancer, such as wasting of adipose and muscle tissues, gut atrophy, microenvironmental extracellular matrix remodeling and immune cell infiltration [21–24]. Although the discovery and functions of the Toll-NF-κB and IMD-NF-κB pathways were pioneered in flies and found to control developmental patterning and infection response of the innate immune system [25–27], less information is available for its possible roles in carcinogenesis.

Here, we describe the aberrant expression and function of a NF-kB transcription factors in the *Ras^V12^; scrib^-^*tumour model. We demonstrate that in *Ras^V12^; scrib^IR^*tumours, the NF-kB transcription factor Dorsal (Dl) plays a pro-tumorigenic role by opposing differentiation and promoting survival and invasion. In this context, Dorsal can amplify JNK signaling activation, a known driver of invasion in this model. We also found that the known Dorsal targets *snail* and *twist* contribute to tumour growth. Interestingly, we report the expression of two different splicing variants of Dl, DorsalA (DlA) and DorsalB (DlB), in non-overlapping tumour cell populations.

## Results

### The Toll pathway promotes tumour growth in an autonomous manner

To identify potential tumour-driving mechanisms, we surveyed upregulated genes in *Ras^V12^; scrib^-/-^* malignant tumours by RNA Sequencing (Fig 1A). We identified 364 genes with elevated expression levels. These include 14 previously identified genes with defined functions in driving *Drosophila* tumor growth or paraneoplastic effects, like organ wasting and diuretic dysfunction (Table 1). Notably, among the upregulated genes - PGRP-SA, GNBP2, Easter, Spz3, and Spz5 - encode extracellular components of the Toll pathway (Fig 1A). The Toll pathway, one of the two established NF-kB signaling pathways in *Drosophila*, is differentially activated during development and in response to fungal or bacterial infection. PGRP-SA (Peptidoglycan Recognition Protein SA) is a secreted protein that binds to bacteria-derived peptidoglycans upon bacterial infection, forming a trimeric complex with a member of the GNBP family (Gram-Negative Binding Protein) and the serine protease ModSP (Modular Serine Protease). This association initiates a multi-step proteolytic cascade, eventually cleaving the ligand Spz (Spaetzle). Subsequently Spz binds to the Toll receptor, activating the Toll pathway intracellularly (Fig 1B). To date, six Spz ligands and nine Toll receptors have been identified and the functions and associations of the different ligands and receptors is not yet fully understood [27]. Intracellularly, Toll receptor activation triggers the activation and formation of a complex composed of the adaptor proteins Myd88 and Tube as well as the serine-threonine protein kinase Pelle. Pelle subsequently phosphorylates the inhibitor Cactus, the homolog of mammalian IkappaB, which is targeted to proteasome degradation. As Cactus sequesters the NF-kB Dorsal or Dif into the cytoplasm, its degradation triggers the translocation of the NF-kBs into the nucleus and the transcription of their target genes (Fig 1B).

**Fig 1.**
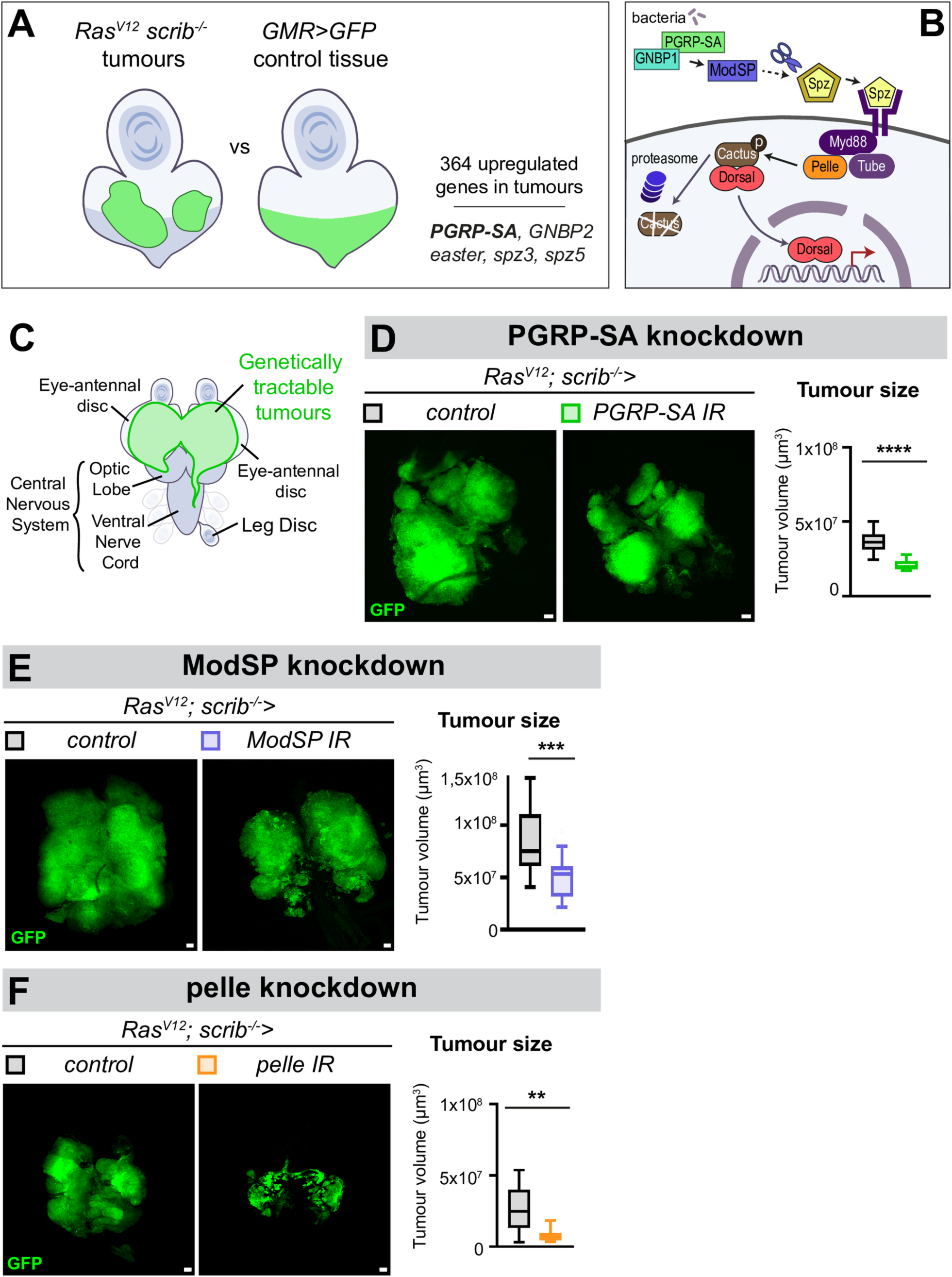
Extracellular and intracellular Toll signaling components contributes to *Ras^V12^; scrib^-/-^*tumour growth. (A) A transcriptomics analysis comparing the profile of *Ras^V12^; scrib^-/-^* tumour cells to *GMR>GFP* control cells in the eye-antennal disc (EAD) identified 364 genes that are upregulated in the tumours. Among them, *PGRP-SA*, *GNBP2*, *easter*, *spz3* and *spz5* are genes taking part in the extracellular machinery leading to Toll pathway activation. (B) Simplified version of Toll pathway activation upon bacterial infection. PGRP-SA (Peptidoglycan recognition protein SA) is a carboxypeptidase which recognizes and binds to peptidoglycans upon bacterial infection. Follows the formation of a trimeric complex together with a member of the GNBP family (Gram Negative Binding Protein) and the serine protease (ModSP). This complex initiates a multi-step proteolytic cascade that ultimately leads to the cleavage and activation of the ligand Spz. After the binding of Spz to the Toll receptor, the pathway is activated intracellularly through the activation and formation of a complex composed of the adaptor proteins Myd88 and Tube as well as the serine-threonine protein kinase Pelle. Pelle subsequently phosphorylates the inhibitor cactus, the homolog of mammalian IkappaB, which is targeted to proteasome degradation. As Cactus sequesters the NF-kB Dorsal into the cytoplasm, its degradation triggers the translocation of Dorsal into the nucleus and the transcription of its target genes. (C) Cartoon of the genetic setting used in this study for manipulating *Ras^V12^; scrib^-/-^*tumours. GFP-labelled tumours are generated from randomly-selected single epithelial cells from the EAD. They grow and fuse to form large tumours that start invading the central nervous system (CNS) through the Optic Lobe (OL) first and the Ventral Nerve Cord (VNC) later. Tumour cells also sometimes reach the leg discs (LD). (D) Representative confocal pictures and quantifications of the mean tumour volumes of GFP-labelled control tumours (n=21, m=3,71x10^7^ µm^3^, SD=±0,47x10^7^ µm^3^) and *PGRP-SA Ras^V12^; scrib^-/-^* tumours (n=17, m=2,11x10^7^ µm^3^, SD=±0,19x10^7^ µm^3^) at day8 after egg laying, statistical significance was determined with an unpaired T-test with Welch’s correction. (E) Representative confocal pictures and quantifications of the mean tumour volumes of GFP-labelled control (n=20, m=8,46x10^7^ µm^3^, SD=±1,77,53x10^7^ µm^3^) and *ModSP^IR^ Ras^V12^; scrib^-/-^ Dcr2* tumours (n=30, m=5,07x10^7^ µm^3^, SD=±1,00x10^7^ µm^3^) at Day8 (29°C), statistical significance was determined with an unpaired T-test with Welch’s correction. (F) Representative confocal pictures and quantifications of the mean tumour volumes of GFP-labelled control tumours (n=10, m=2,64x10^7^ µm^3^, SD=±0,76x10^7^ µm^3^) and *pelle^IR^ Ras^V12^; scrib^-/-^* tumours (n=10, m=0,78x10^7^ µm^3^, SD=±0,22x10^7^ µm^3^) at Day6 (29°C), statistical significance was determined with a Mann-Whitney test. Scale bars=50µm.

**Table 1.**
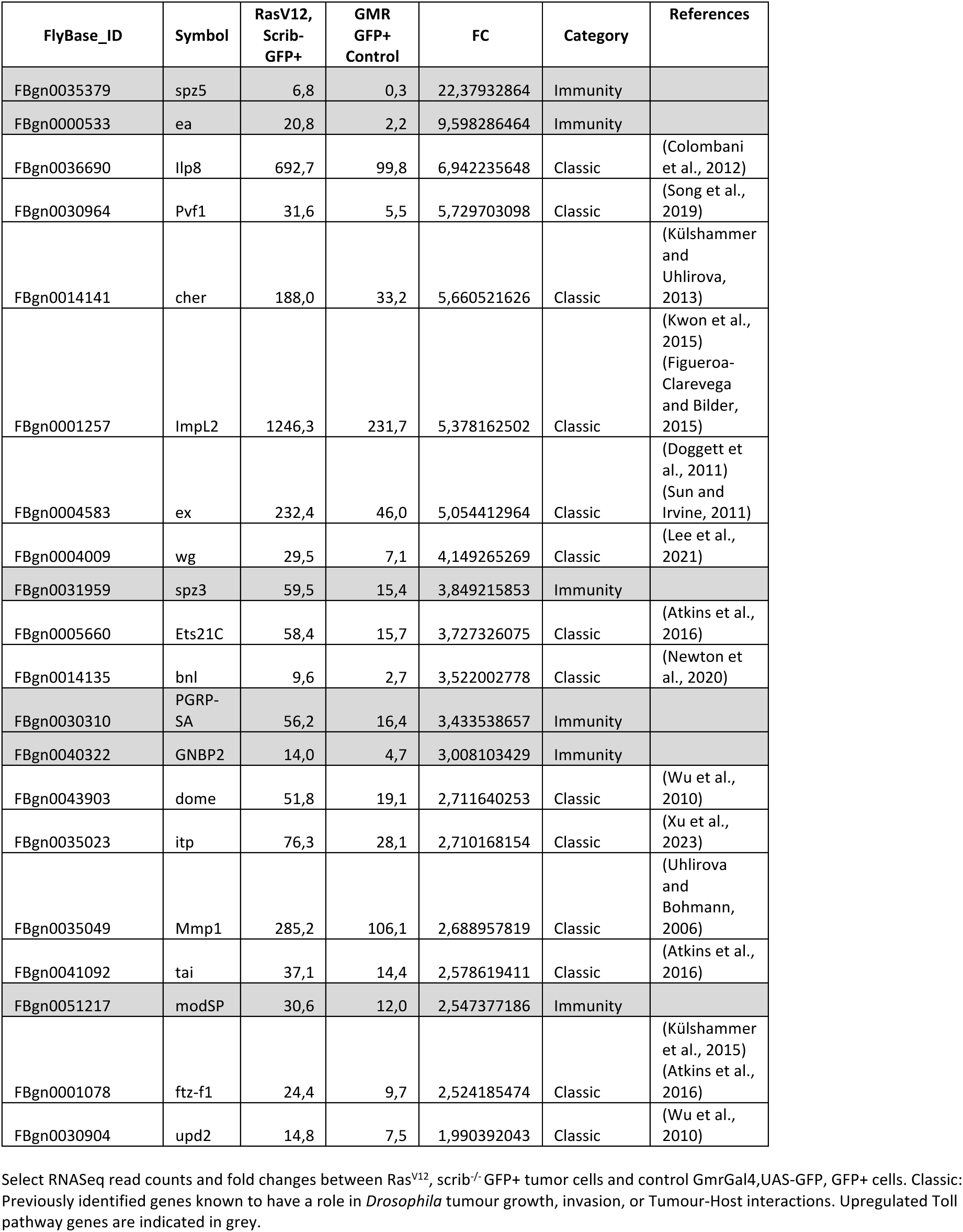
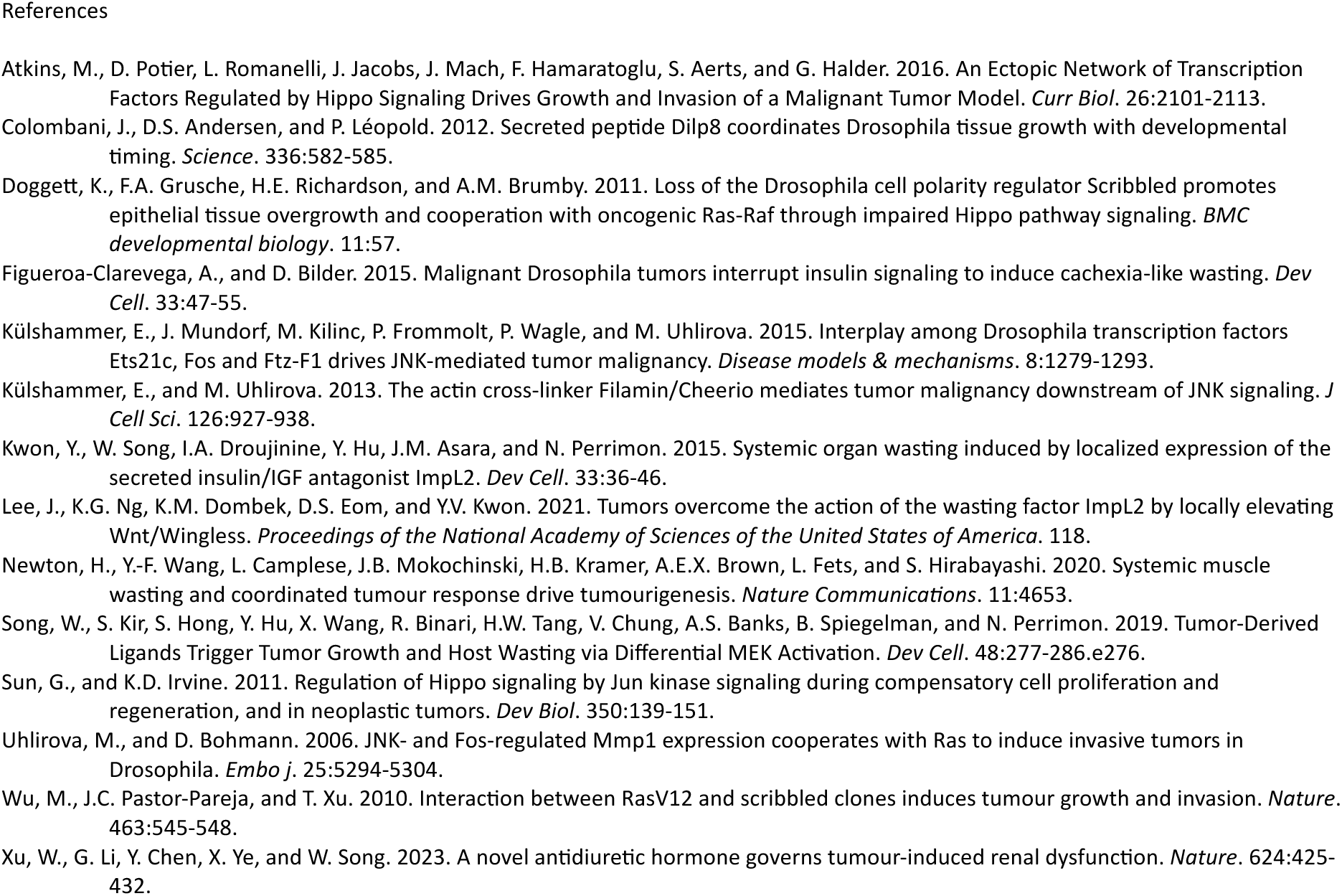
Select list of genes previously reported to have a role in *Drosophila* tumor models alongside Toll pathway components.

We decided to first investigate whether PGRP-SA is involved in *Ras^V12^; scrib^-/-^* tumorigenesis. To do so, we genetically induced the formation of *Ras^V12^; scrib^-/-^* tumours from randomly induced single cells in the Eye-Antennal Disc (EAD) of the *Drosophila* larva. This way, we can knockdown or overexpress any gene of interest within the tumour specifically without affecting expression in the rest of the body (Fig 1C). To improve RNA interference (IR) expression levels and processing respectively, knockdown experiments were sometimes performed at 29°C with the additional expression of Dicer-2, as indicated in the legends.

Knock-down of PGRP-SA (*PGRP-SA^IR^*) significantly reduced *Ras^V12^; scrib^-/-^* tumour growth compared to control tumours (Fig 1D). Similarly, knock-down of ModSP (*ModSP^IR^*) significantly decreased tumour size (Fig 1E), suggesting that the extracellular components of the Toll pathway surprisingly play a pro-tumorigenic role in this context. To address whether Toll signaling is required within tumour cells or in another microenvironmental compartment, we performed knock-down of the intracellular kinase Pelle/IRAK (*pelle^IR^*), a hub in Toll signal transduction. *pelle^IR^ Ras^V12^; scrib^-/-^* tumours exhibited a drastic size reduction compared to control tumours (Fig 1F). These results collectively suggest that *Ras^V12^; scrib^-/-^* tumours produce extracellular Toll-activating components that contribute to a Toll pathway pro-tumorigenic signal activity in an autonomous manner.

### Dorsal is ectopically expressed in oncogenic *Ras^V12^*-driven tumours

Toll pathway activation leads to the release and nuclear translocation of the NF-κB transcription factors Dl and/or Dif, leading to transcription of target genes. While most studies of Dl and Dif in developmental patterning and infection have focused on their main splicing variants (DlA and DifA), both NF-kB transcription factors have been reported to also have alternatively spliced B isoforms (DlB and DifB) [28, 29]. For instance, both DlA (678 amino acids) and DlB (994 amino acids) contain a Relish-homology domain, composed of a DNA binding domain and a dimerization domain (for homo/heterodimerization and Cactus binding), as well as a transactivation domain (for transcription activation) [28]. However, the transactivation domain of DlA is much shorter than Dorsal-B. Noteworthy, only DlA possesses an NLS sequence. Therefore, DlB cannot translocate to the nucleus by itself but is thought to be able to bind to DlA for translocating to the nucleus of fat body cells upon infection [28]. Contrary to DlA, DlB is not expressed and involved in embryonic dorso-ventral patterning. It is rather specifically expressed in the subsynaptic reticulum of neuromuscular junctions, where it stabilizes Cactus. Similarly, Dif-B is found in the mushroom body of the CNS, where it also stabilizes Cactus [29]. Although these observations support the idea that B isoforms play a specific role in nervous system tissues, their precise functions remain to be elucidated [30].

We began by assessing the expression of Dl in *Ras^V12^; scrib^IR^* tumours through immunostainings against DlA and DlB (Fig 2A, 2B, 2C, 2D and S1A, S1B Fig). Both Dorsal isoforms were highly expressed in tumors in the posterior part of the disc that attaches to the brain through the prospective optic stalk (Fig 2A and 2B), while neither DlA or DlB proteins were observed in control clones (Fig 2A). DlA and DlB were almost entirely mutually exclusive, in different cell populations (Fig2A, 2B and S1B Fig). Within *Ras^V12^; scrib^IR^* tumours, cells can be easily separated according to their levels of DlA mean intensity (Fig 2C). We identified a DlA^high^ cell population that had four-fold higher DlA protein levels compared to the DlA^low^ remaining cell population. Importantly, the DlA mean intensity in the DlA^low^ cell population was consistently higher than in the surrounding *wt* epithelial cells of the EAD (Fig 2C). These observations suggest that although we observed a peak of DlA expression at the posterior part of the tumour, there is a general elevation of the basal level of DlA throughout the entire tumour. In contrast, we did not observe a general elevation of DlB throughout the tumour when compared to *wt* cells of the TME, but we did observe a local increase in the posterior part of the tumour of a similar magnitude to DlA (Fig 2B and 2D). Tumour cells located in the posterior area of the tumour are closest to the CNS, which is the first organ these tumours invade [31–33]. Intriguingly, tumour cells that have initiated invasion into the optic lobes (OL) of the CNS displayed high levels of DlA or DlB (S1B Fig), suggestive of a potential role of Toll-NFκB signaling in tumor cell migration.

**Fig 2.**
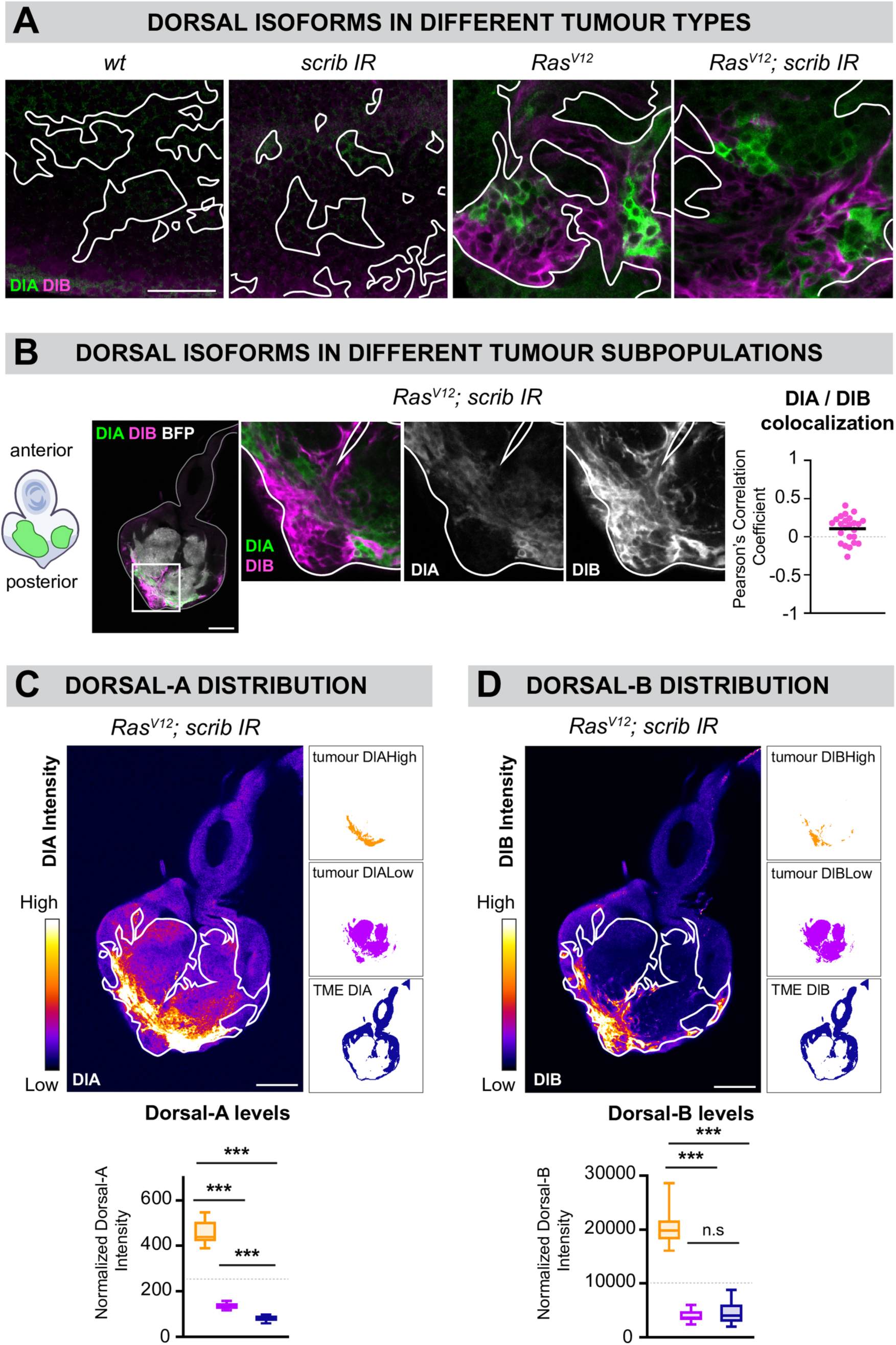
Two splicing variants of the NF-kB Dorsal are aberrantly expressed in *Ras^V12^*-driven tumours. (A) Immunostainings against Dorsal-A (DlA in green) and Dorsal-B (DlB in magenta) in BFP-labelled *wt* clones (n=20 EADs), *scrib^IR^* tumours (n=23 EADs), *Ras^V12^* tumours (n=20 EADs) and *Ras^V12^; scrib^IR^*tumours (n=24 EADs) at day 6. DlA and DlB accumulation is observed only in *Ras^V12^*-driven tumours and appears mainly mutually exclusive. (B) Immunostaining against DlA (green) and DlB (magenta) in BFP-labelled *Ras^V12^; scrib^IR^* tumours (white) (n=24 EADs) and close-ups showing that DlA and DlB are not expressed in the same subset of cells. Colocalization analysis assessed with the Pearson’s correlation coefficient reveals a poor colocalization between DlA and DlB in the tumours (n=24, m=0,11, SD=±0,08). (C) Heat map of DlA intensity within a representative BFP-labelled *Ras^V12^; scrib^IR^* tumour-bearing EAD at day 6. The tumour is outlined with the straight white line. An arbitrary intensity threshold set at 250 (gray line on the graph) allows the clear segregation of a DlAhigh (“tumour High”) and a DlAlow (“tumour Low”) cell population within the tumour. Quantifications of the mean DlA intensity of the “tumour High” (n=23, m=458,7, SD=±28,4), “tumour Low” (n=23, m=135,2, SD=±7,0) and wt Tumour microenvironment (TME) (n=23, m=82,2, SD=±6,7) demonstrates a general elevation of DlA levels within the tumour compared to wt cells. Statistical significance was determined with a Tukey’s multiple comparison test. (D) Heat map of DlB intensity within a representative RasV12; scribIR tumour-bearing EAD at day 6. The tumour is outlined with the straight white line. An arbitrary intensity threshold set at 10000 (gray line on the graph) allows the clear segregation of a DlBhigh (“tumour High”) and a DlBlow (“tumour Low”) cell population within the tumour. Quantifications of the mean DlB intensity of the “tumour High” (n=24, m=20331,9, SD=±3108,7), “tumour Low” (n=24, m=3955,8, SD=±584,5) and wt Tumour microenvironment (TME) (n=23, m=4660,9, SD=±1187,3). Normalization was done by subtracting the background intensity. Statistical significance was determined with a Dunn’s multiple comparison test. Scale bars=20µm.

Finally, a strong enrichment of the repressor Cactus, the ortholog of mammalian IkappaB (IKKB), was observed in DlA^high^ tumour cells but not in DlB^high^ tumour cells, as assessed with an endogenous GFP fusion protein reporter, *Cactus::GFP* (S1C Fig). This is particularly interesting as *cactus* is a positive transcriptional target of the Toll pathway[34]. Cactus upregulation constitutes a negative feedback loop which could cap Toll pathway activation, a mechanism that has also been described in mammalian NF-kB signalling pathways [35, 36]. In conclusion, simultaneous elevation of DlA, DlB and Cactus expression suggests that in the posterior part of *Ras^V12^; scrib^IR^* tumours, Toll pathway activation is significantly higher than in the rest of the tumour.

### Dorsal aberrant accumulation is genetically and spatiotemporally-regulated

We investigated what could be the upstream mechanism leading to aberrant Dl accumulation. Initially, we examined Dl isoforms levels in various genetic backgrounds: control *wild type* clones, *scrib^IR^* tumours, *Ras^V12^* tumours and *Ras^V12^; scrib^IR^* tumours. We observed aberrant expression of DlA as well as DlB exclusively in *Ras^V12^*and *Ras^V12^; scrib^IR^* tumours (Fig 2A), suggesting that *Ras^V12^* instructs tumour cells competent to produce these isoforms at abnormal levels. Furthermore, we followed DlA levels over time in *Ras^V12^; scrib^IR^* tumours, using an endogenous Dorsal genetic reporter, that acts as a Dorsal-A specific reporter in this context (Fig 3A and S2A and S2B Fig). The onset of DlA expression correlated with the appearance of Elav-positive neurons in the prospective eye region of the tumours around Day5 (Fig 3A). Interestingly, DlA accumulate in cells surrounding the neurons but was rarely detected in the neurons themselves (Fig 3A yellow arrows and Fig 3B). Concurrently, DlB expression was exclusive to the tumour neurons (Fig 3C). Within the tumours, apoptosis was predominantly observed in the differentiating areas where we observed dying neurons (Fig 3D, yellow arrows). Therefore, we hypothesized that the local aberrant Dl expression might be triggered by cell death-derived signals. However, efficient blocking of apoptosis in *Ras^V12^; scrib^IR^* tumours through Diap1 overexpression (*Diap1^OE^*) did not affect increased DlA and DlB levels (Fig 3E and 3F).

**Fig 3.**
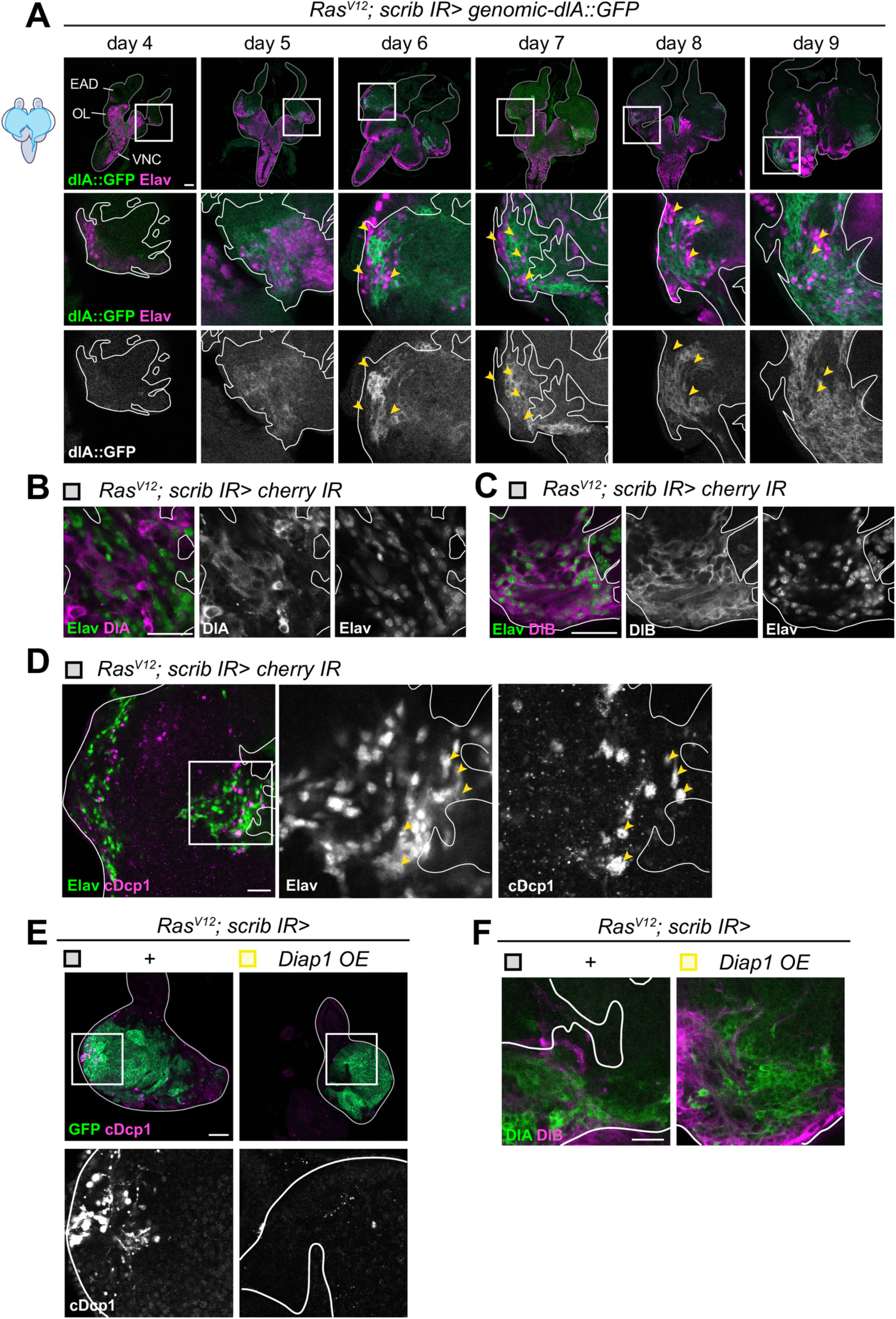
Cell type-dependent Dorsal isoforms accumulation correlates with tumour neurogenesis and apoptosis. (A) Representative confocal pictures of BFP-labelled *Ras^V12^; scrib^IR^* tumours, expressing the a DlA::GFP fusion protein at endogenous levels (green), overtime (Day 4 n=18 EADs, Day 5 n= 14 EADs, Day 6 n=18 EADs, Day 7 n= 17 EADs, Day 8 n=20 EADs and Day 9 n=11 EADs) stained for the pan-neuronal marker Elav (magenta). The cephalic complex is outlined with the straight gray line and the tumour is outlined with the straight white line on the close-ups. Yellow arrows points at neurons that are negative for DlA. Scale bar=50µm. (B) Representative confocal pictures of GFP-labelled *cherry^IR^ Ras^V12^; scrib^IR^*(n=8 EADs) at day 6 (29°C) stained for Elav (green) and DlA (magenta). The tumour is outlined with the straight white line. DlA and Elav appears mutually exclusive. Scale bar=20µm. (C) Representative confocal pictures of GFP-labelled *cherry^IR^ Ras^V12^; scrib^IR^* (n=8 EADs) at day 6 (29°C) stained for Elav (green) and DlB (magenta). The tumour is outlined with the straight white line. DlB and Elav are expressed in the same tumour cells. Scale bar=20µm. (D) Representative confocal pictures of GFP-labelled *cherry^IR^ Ras^V12^; scrib^IR^* control tumours (n=62 EADs) at day 6 (29°C) stained for Elav (green) and cDcp1 (magenta). The tumour is outlined with the straight white line. cDcp1^+^ cells are mostly found in areas of the tumour where neurons are present where we often observe neuron apoptosis (yellow arrows). Scale bar=20µm. (E) Representative confocal pictures of GFP-labelled *Ras^V12^; scrib^IR^*control tumours (n=7 EADs) and *Diap1^OE^ Ras^V12^; scrib^IR^* (n=9 EADs) stained for cDcp1 (magenta) at day 6. Scale bar=50µm. (F) Representative confocal pictures of GFP-labelled *Ras^V12^; scrib^IR^*control tumours (n=13 EADs) and *Diap1^OE^ Ras^V12^; scrib^IR^* (n=7 EADs) stained for DlA (green) and DlB (magenta) at day 6. Scale bar=20µm.

Together, these observations demonstrate that the aberrant expression NF-kB/Dl in *Ras^V12^; scrib^IR^* tumours is both genetically and locally controlled. General elevated levels of DlA in the whole tumor population suggests a transcriptional response downstream of oncogenic Ras^V12^ signaling. On top of this Ras^V12^-dependent genetic control, a temporal and or local control triggers elevated levels of DlA, DlB and Cactus specifically in the posterior part of the tumour in the prospective eye region, as well as in invading tumour cells to the brain.

### Dorsal represses Ras^V12^ scrib^IR^ tumour differentiation and death

To investigate the primary functions of Dl in *Ras^V12^; scrib^IR^* tumours, we conducted knockdown experiments using two independent RNAis that efficiently target both Dorsal isoforms (S3A and S3B Fig). *Dorsal^IR^, Ras^V12^; scrib^IR^* tumours exhibited a significant reduction in size compared to *Cherry^IR^ Ras^V12^; scrib^IR^* control tumours (Fig 4A), mimicking the phenotypes observed upon *PGRP-SA*, *ModSP* and *pelle* knockdowns (Fig1C, 1D and 1E). *Ras^V12^; scrib^IR^* tumour growth results from a combination of cell growth, cell proliferation, inhibition of the differentiation of epithelial tumour cells into Elav-positive photoneurons and survival [31, 37]. To delineate the potential functions of Dl in *Ras^V12^; scrib^IR^*tumours, we assessed alterations in proliferation, differentiation and apoptosis. Quantification of mitotic cells suggests that *Dorsal^IR^ Ras^V12^; scrib^IR^* tumours display similar proliferation rates to control tumours, but both their differentiation and apoptosis rates were significantly higher than in control tumours (Fig4B, 4C and 4D). Although, a few neurons form in *Ras^V12^; scrib^IR^* tumours, they appeared spaced apart and did not form the full complement of eight neurons per ommatidia. However, upon Dorsal knockdown, Elav-positive cells increased in number and density, suggesting a partial rescue of differentiation (Fig4C). The fact that we frequently observed cell death coinciding with areas where Elav-positive neurons were present (Fig 3D) suggests that cell death might be a consequence of aberrant tumour cell differentiation. Importantly, *Pelle^IR^ Ras^V12^; scrib^-/-^* tumours also exhibited increased apoptosis and differentiation rates (S3C Fig). This emphasizes the need for the Toll signalling pathway, and not an alternative way to activate Dorsal/NFkB activity, to promote tumour growth. Thus, the Toll-Dl signaling pathway promotes tumour growth primarily by preventing differentiation and promoting survival.

**Fig 4.**
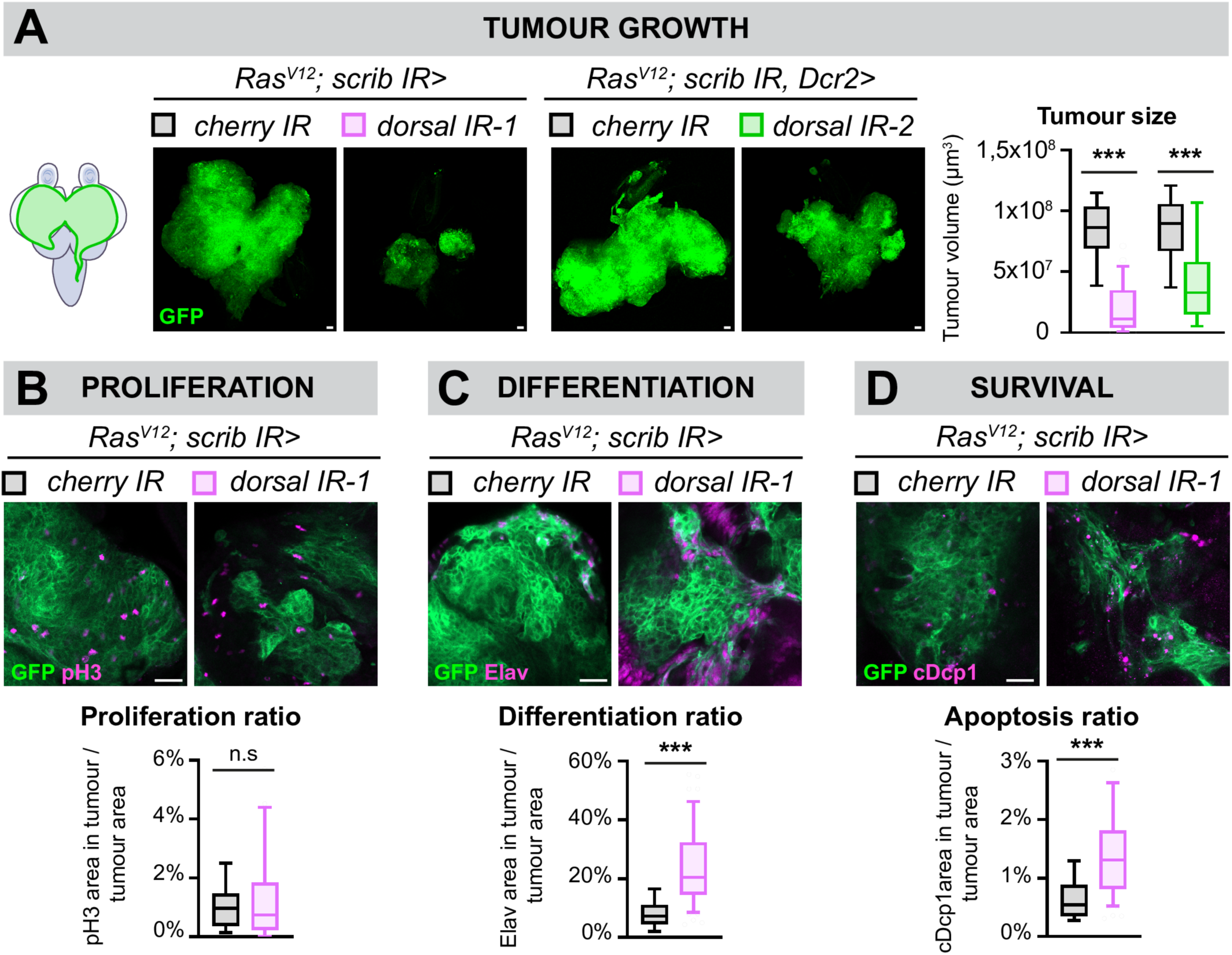
Dorsal represses *Ras^V12^ scrib^RNAi^* tumour differentiation and apoptosis. (A) Representative confocal pictures and quantifications of the mean tumour volumes of GFP-labelled *Cherry^IR^ Ras^V12^; scrib^IR^*control tumours (n=36, m=8,32x10^7^µm^3^, SD=±1,31 x10^7^µm^3^), *dorsal^IR-1^ Ras^V12^; scrib^IR^* tumours (n=25, m=1,97x10^7^µm^3^, SD=± 1,00x10^7^µm^3^), *Cherry^IR^ Ras^V12^; scrib^IR^ Dcr2* control tumours (n=43, m=8,52x10^7^µm^3^, SD=±1,64x10^7^µm^3^) and *dorsal^IR-2^ Ras^V12^; scrib^IR^ Dcr2* tumours (n=37, m=4,71x10^7^µm^3^, SD=±1,85x10^7^µm^3^) at Day12 (29°C), statistical significance was determined with an unpaired T-test with Welch’s correction. Scale bar=50µm. (B) Representative confocal pictures and quantifications of the proliferation ratio of GFP-labelled *Cherry^IR^ Ras^V12^; scrib^IR^* control tumours (n=36, m=1,08%, SD=±0,41%) and *dorsal^IR-1^ Ras^V12^; scrib^IR^* tumours (n=35, m=1,39%, SD=±0,83%) at Day8 (29°C). Proliferation was detected through immunostainings against the mitotic marker pH3 (phospho-Histone 3, in magenta). Here, the proliferation ratio is defined as the ratio between the pH3 area within the tumour and the area of the tumour itself. Scale bar=20µm. (C) Representative confocal pictures and quantifications of the differentiation ratio of GFP-labelled *Cherry^IR^ Ras^V12^; scrib^IR^* control tumours (n=63, m=8,29%, SD=±2,87%) and *dorsal^IR-1^ Ras^V12^; scrib^IR^* tumours (n=57, m=24,40%, SD=±7,09%) at Day8 (29°C). Differentiation was detected through immunostainings against the pan-neuronal marker Elav (magenta). Here, the differentiation ratio is defined as the ratio between the Elav area within the tumour and the area of the tumour itself. Scale bar=20µm. (D) Representative confocal pictures and quantifications of the apoptosis ratio of GFP-labelled *Cherry^IR^ Ras^V12^; scrib^IR^* control tumours (n=63, m=0,71%, SD=±0,29%) and *dorsal^IR-1^ Ras^V12^; scrib^IR^* tumours (n=57, m=1,45%, SD=±0,46%) at Day8 (29°C). Apoptosis was detected through immunostainings against the cleaved effector caspase, cDcp1 (cleaved-Death Caspase 1, in magenta). Here, the apoptosis ratio is defined as the ratio between the cDcp1 area within the tumour and the area of the tumour itself. Statistical significance was determined with an unpaired T-test with Welch’s correction. Scale bar=20µm.

### Dorsal is required for invasion and accentuates JNK signaling

We next addressed whether Dl is involved in tumor invasion. In the *Ras^V12^; scrib^IR^* tumour model, cells adopt a migratory behavior by JNK signaling. Specifically, JNK mediates expression of Matrix MetalloProteinase 1 (MMP1) and Cheerio (Cher-mammalian Filamin), resulting in basement membrane degradation and cytoskeleton remodeling respectively [22, 32, 37–41]. We conducted colocalization analyses, assessed by the Pearson’s Correlation Coefficient (PCC), between the Dorsal isoforms and JNK activity in the tumours, assessed either through activated JNK (phosphor-JNK, pJNK) or a JNK activity reporter, TRE-eGFP. Overall, we found a low colocalization between DlA and TRE-eGFP (Fig 5A) while we observed a good PCC between DlB and pJNK within the tumours (Fig 5B). These observations align with the finding that tumor cells invading the CNS often have high levels of either DlA or DlB.

**Fig 5.**
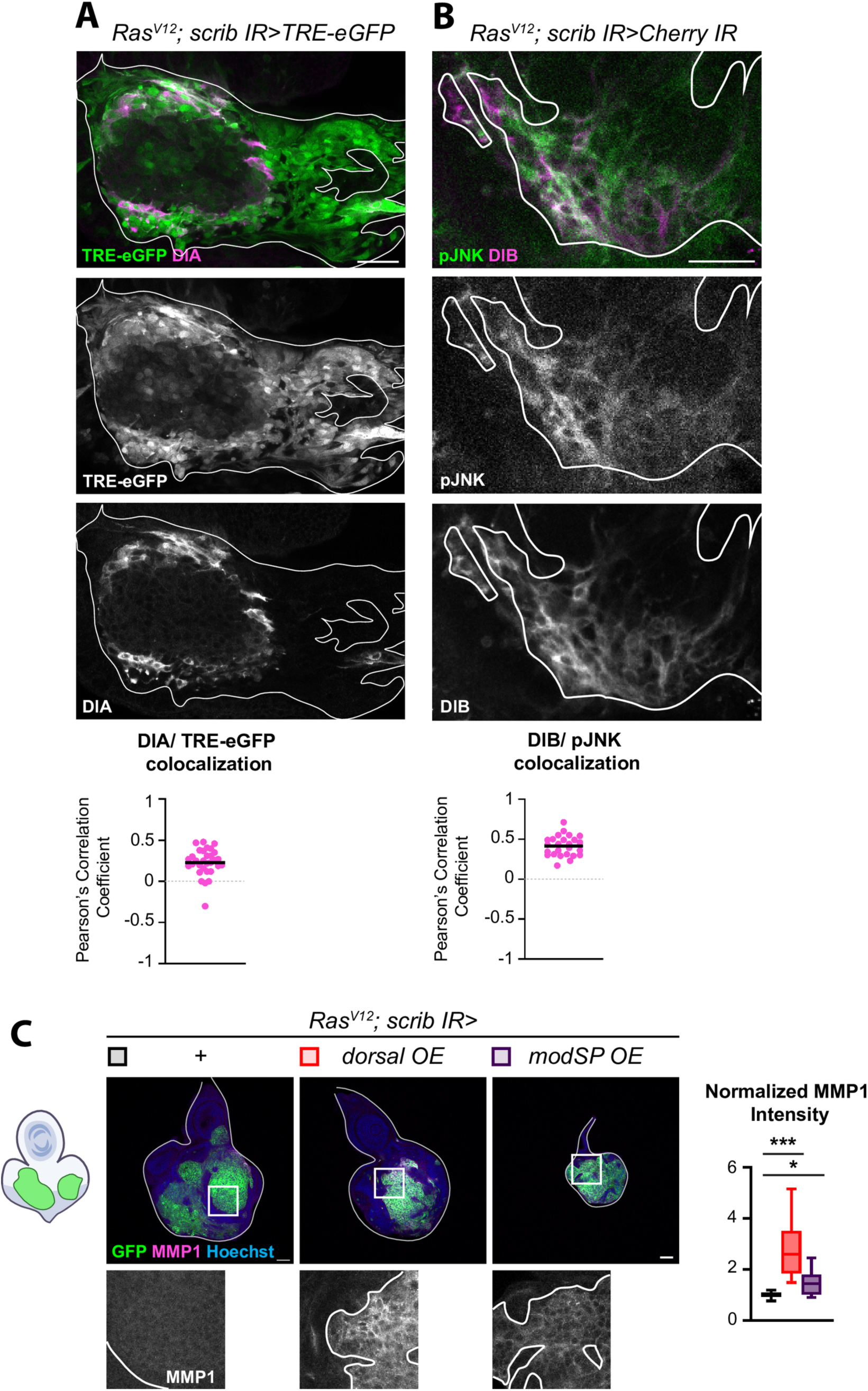
Dorsal amplifies JNK signaling. (A) Representative confocal pictures of BFP-labelled *Ras^V12^; scrib^IR^* tumours, expressing the JNK activity reporter, TRE-eGFP (green) and stained for DlA (magenta). The tumour is outlined with the straight white line. Colocalization analysis assessed with the Pearson’s correlation coefficient reveals a weak colocalization between DlA and JNK activity in the tumours (n=31, m=0,23, SD=±0,08). Scale bar=20µm. (B) Representative confocal pictures of GFP-labelled *cherry^IR^ Ras^V12^; scrib^IR^* tumours stained for pJNK (green) and DlB (magenta) at day6 (29°C). The tumour is outlined with the straight white line. Colocalization analysis assessed with the Pearson’s correlation coefficient reveals a good colocalization between DlB and pJNK in the tumours (n=25, m=0,41, SD=±0,06). Scale bar=20µm. (C) Representative confocal pictures and quantifications of the mean MMP1 intensity of GFP-labelled *Ras^V12^; scrib^IR^* tumours (n=11, m=1,00, SD=±0,06), *dorsal^OE^ Ras^V12^; scrib^IR^* tumours (n=11, m=2,85, SD=0,57) and *modSP^OE^ Ras^V12^; scrib^IR^* tumours (n=10, m=1,49, SD=±0,32) at day 6. Intensity normalization was done by subtracting the background intensity on each picture and finally dividing each data point by the mean intensity value for the control tumours for each replicate. Statistical significance was determined with Dunnett’s T3 multiple comparisons test. Scale bars=50µm.

To test whether Dorsal may function upstream of JNK signaling in *Ras^V12^; scrib^IR^* tumours, we assessed the levels of the JNK target MMP1 following *Dorsal^OE^* and *ModSP^OE^* overexpression which enhances Toll signaling and subsequently Dorsal activity. MMP1 mean intensity was significantly higher in tumors with elevated *Dorsal* or *ModSP* expression. compared to control tumours. Therefore, Toll-Dl signaling can augment expression of the JNK transcriptional target, *mmp1* in *Ras^V12^; scrib^IR^* tumours (Fig 5C).

As both DlA, DlB and Cactus are highly expressed in brain lobe proximal and invading cells and Dl amplifies JNK signalling in the tumours, we investigated the effect of Dorsal knock-down on tumour cell migration. We scored for three parameters reflecting the mobility of tumour cells at late stages of tumour development (Day 12): 1/ the VNC invasion score, which assigns a higher score to tumors with greater migration toward the tip of the ventral nerve cord (VNC) of the CNS, 2/the frequency at which we observe tumour cells within the organs adjacent to the VNC called leg discs and 3/ the fusion of tumours generated in the two distinct EAD, indicating the ease with which cells move from one EAD to the other and EADs and come into contact with each other (Fig 6A). To mitigate the influence of tumour cell number on migration assessment, we compared control and *Dorsal^IR^* tumours of similar size. Upon Dorsal knock-down, at late stages, the three parameters showed a drastic decrease: the VNC score decreased threefold compared to control, reaching a low value similar to early-stage control tumours (Fig6B). The frequency of invasion into leg discs decreased sevenfold and tumors from neighboring EADs often failed to merge (Fig 6A). Similarly, Dorsal overexpression (*Dorsal^OE^*) increased tumour cell mobility even at early stages (Day 6), while tumour size is signicantly smaller (Fig 6B), suggesting that 1/ the increase in invasion is not simply a secondary consequence of higher cell number and 2/ that the balance between invasion and tumour growth tilted toward invasion when tumour cells have high levels of Dorsal.

**Fig 6.**
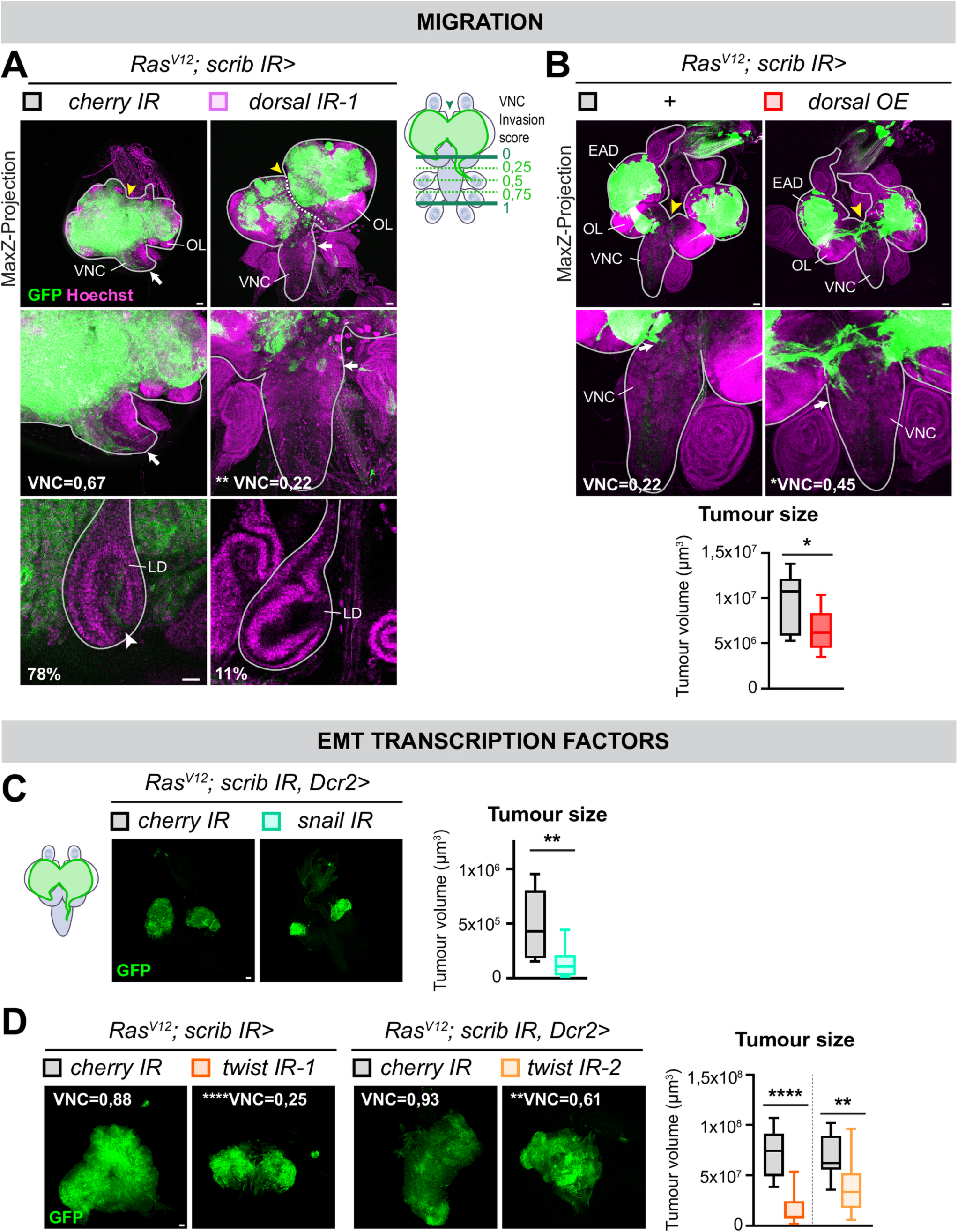
Dorsal stimulates tumour migration and the Dorsal targets Snail and Twist contribute to tumour growth. (A) Z-projections of representative confocal pictures of size-matched GFP-labelled *Cherry^IR^ Ras^V12^; scrib^IR^* tumours (n=3 EADs) and *dorsal^IR-1^ Ras^V12^; scrib^IR^* tumours (n=8 EADs) at day 12 (29°C), stained for Hoechst (magenta). Scale bars=50µm. The cephalic complexes (and leg discs on the small inserts) are outlined with the straight gray line. A cartoon represents the three invasion parameters assessed in this study: the VNC (Ventral Nerve Cord) invasion score, ranged between c and 1, is greater the further tumour cells migrate toward the tip of the VNC (green arrow), indicated as “VNC=” on the Z-projection images; the frequency of leg discs invasion (small insert with leg discs confocal pictures, Scale bars=20µm) and the fusion of distinct EAD (Eye-Antennal Disc) tumours (white arrow, dashed line indicates the visible separation of both neighbor EADs). For the VNC score, statistical significance was determined with Dunnett’s T3 multiple comparisons test. (B) Z-projections of representative confocal pictures of GFP-labelled and *Ras^V12^; scrib^IR^* (n=15 EADs) and *dorsal^OE^ Ras^V12^; scrib^IR^* tumours (n=24 EADs) at day 6, stained for Hoechst (magenta). The cephalic complexes (and leg discs on the small inserts) are outlined with the straight gray line. Quantification of the mean tumour volumes of GFP-labelled *Ras^V12^; scrib^IR^* (n=9, m=9,71x10^6^µm^3^, SD=±1,53x10^6^µm^3^) and *dorsal^OE^ Ras^V12^; scrib^IR^* tumours (n=11, m=6,52x10^6^µm^3^, SD=±1,12x10^6^µm^3^) at day 6. OL: Optic Lobe. Statistical significance was determined with an unpaired T-test with Welch’s correction. Scale bars=50µm. (C) Representative confocal pictures and quantification the mean tumour volume of GFP-labelled *cherry^IR^ Ras^V12^; scrib^IR^* (n=12, m=4,83x10^5^µm^3^, SD=±1,52x10^5^µm^3^), and *snail^IR^ Ras^V12^; scrib^IR^* (n=9, m=1,41x10^5^µm^3^, SD=±0,65 x10^5^µm^3^). Statistical significance was determined with a Mann Whitney’s test. (D) Representative confocal pictures and quantification the mean tumour volume of GFP-labelled *cherry^IR^ Ras^V12^; scrib^IR^*(n=20, m=7,17x10^7^µm^3^, SD=±1,22x10^7^µm^3^) and *twist^IR-1^ Ras^V12^; scrib^IR^* (n=7, m=1,81x10^7^µm^3^, SD=±0,83x10^7^µm^3^) as well as *Cherry^IR^ Ras^V12^; scrib^IR^ Dcr2* control tumours (n=24, m=6,89x10^7^µm^3^, SD=±1,12x10^7^µm^3^) and twist*^IR-2^ Ras^V12^; scrib^IR^ Dcr2* tumours (n=13, m=3,98x10^7^µm^3^, SD=±1,43x10^7^µm^3^). VNC invasion scores are indicated. Statistical significance was determined with a Mann Whitney’s test. Scale bars=50µm.

Given the functions of Dorsal in the tumour context, we wondered whether *Dorsal^OE^* is sufficient to induce tumorigenesis. Although, *Dorsal^OE^* control clones were not bigger than *wt* clones, we systematically observed the formation of dysplasic GFP-labeled *Dorsal^OE^* clones within the EADs that appeared as large-rounded cell masses often located basally (S4A and S4B Fig, yellow arrow). We hypothetize that they arise from the inhibition of differentiation. Moreover, in this context, we never observed *Dorsal^OE^* cells in the VNC (VNC score in S4A Fig). Thus, Dorsal at high levels is sufficient to trigger dysplasia but not invasive hyperplastic tumours.

The Epithelial-to-Mesenchymal (EMT) transcription factor Snail has been implicated in promoting invasion of *Ras^V12^; lgl^-/-^*tumours (a similar tumour model to *Ras^V12^; scrib^-/-^*tumours) through stimulation of JNK signaling and EMT-like cytoskeleton remodeling [42]. Snail, along with Twist, another EMT transcription factor, are well-known targets of Dorsal that govern mesoderm invagination during developmental patterning in cells with the highest Dorsal activity [43–46]. Interestingly, knockdowns of both Snail and Twist significantly decreased *Ras^V12^; scrib^IR^*tumour size (S4A and S4B Fig), mimicking Dorsal knockdown. At late stages, twistIR *Ras^V12^; scrib^IR^* tumour also displayed a lower VNC score compared to control tumours, suggesting impaired tumour cell migration (S4B Fig). We propose a model where the Dl^high^ tumour cells activate Twist and Snail expression, provoking migration, in part through JNK signalling enhancement (Fig 7A).

**Fig 7.**
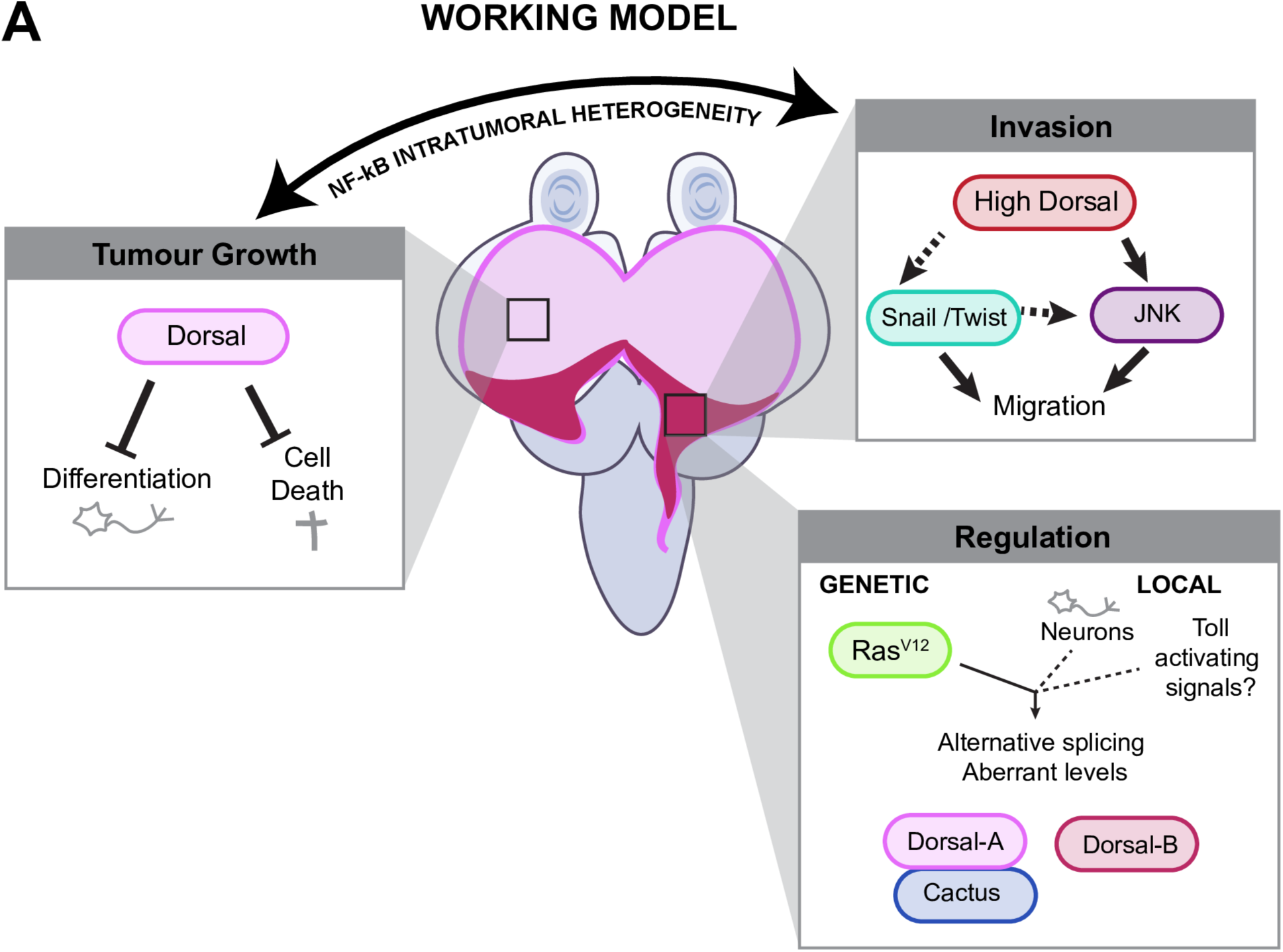
Working Model. In this working model, we propose that the protein level of the NF-kB Dorsal regulate different aspect of tumorigenesis. Invasion: In the most posterior part of the tumour, high levels of Dorsal promote invasion of the tumour cells in the adjacent organs through elevation of JNK signaling and the EMT transcription factor Snail and Twist that would promote cytoskeleton remodelling. Tumour growth: In the rest of the tumour where more intermediate levels of Dorsal are observed, Dorsal would stimulate tumour growth by repressing differentiation and apoptosis. Regulation: At least two factors seem two trigger aberrant level of Dorsal within the tumours. Genetically, Ras^V12^ make tumour cells competent to accumulate higher levels of Dorsal throughout the entire tumour. Locally, a spatiotemporal control is exerted at the posterior part of the tumour where neurons are formed, to reach very high levels of Dorsal. A possibility is an increase local Toll pathway activation triggered by high Spz concentration from the brain which will promote Dorsal expression. There, we observe alternative splicing of DlA versus DlB which segregate in different tumour subpopulations. We hypothesize that the heterogeneity in Dorsal isoforms and/or levels promote a balance between tumour growth and invasion favoring tumorigenesis.

Taken together, these experiments suggest that Dl promotes tumor growth through repressing cell death and differentiation as well as promoting EMT like delamination and invasion dependent on Twist and Snail activity.

## Conclusion/Discussion

Here, we establish that autonomous activation of the inflammatory Toll pathway plays a pivotal role in supporting the growth of *Ras^V12^; scrib^IR^* tumours. The NF-kB transcription factors Dorsal, and key effectors of the Toll pathway, exhibit aberrant expression in this model. In this context, Dorsal activity facilitates tumour growth by preventing differentiation and promoting survival without measurable impact on proliferation. Additionally, Dorsal promotes invasion genetically dependent on the EMT factors Snail and Twist.

An exciting feature in this tumor model is tumor heterogeneity in both NF-kB protein composition (DlA and DlB) and Toll pathway activity (Dl^high^ cactus^high^ vs Dl^low^, Cactus^low^), as judged by classical target gene expression (*cactus* and *dorsal*) and stimulation of invasion. Dl is unevenly accumulated within the population of transformed cells in a spatial and regional manner and on at least two levels: there is a low-level DlA accumulation within the entire region of Ras^V12^ expression and a heightened level in the posterior region situated in the differentiating prospective eye region that is physically attached to the brain through the stalk. In this region a notable accumulation of both splicing variants of Dl, DlA and DlB, is seen within distinct tumour cell populations.

While NF-kB protein heterogeneity within the same tumour has not been studied yet in solid mammal tumours, the NF-kB fingerprinting and their subsequent homo-/heterodimerization within diffuse large B-cell lymphoma hold predictive values for the tumour cell response to microenvironmental activating cues [47]. With our study, this finding, suggest that NF-kB composition heterogeneity might also occur in solid mammal cancers and modulate their response to microenvironmental inflammation.

Moreover, heterogeneity in the level/activity of a single NF-kB mirrors observations in mammalian cancers. In pancreatic cancer, where oncogenic KRAS is the main driver, tumor heterogeneity is a central hallmark and frequently associated with variable NF-kB signaling. A central question is thus, what causes some cells to respond with inflammation whereas others do not? Possibilities include non-autonomous external input from nearby cells of the microenvironment or circulation versus intrinsic mechanisms. *In vitro* studies of KRAS^G12V^-transfected immortalized human pancreatic epithelial derived cells (HPDE) have demonstrated that variable levels of oncogenic KRAS can produce different outcomes. High KRAS^G12V^ expression can direct NF-kB signaling and EMT phenotypes driven in part by feed forward loop through IL6 signaling, whereas lower level of KRAS activity retains epithelial character with lower level of NF-kB signaling dependent on IL1 [48].

In the *Ras^V12^, scrib^-/-^* model, a “flip out” cassette turns on Ras^V12^ expression under control of the actin promoter in all cells with the same genetic dose, and by first principle should signal evenly. Yet, Ras^V12^ - driven NF-kB expression is temporally and spatially controlled in a stereotypic pattern. One possibiity is that this region is prepatterned to respond to increased levels of Ras^V12^ with a higher response. Indeed, in this region, we observe a higher level of pMAPK and where tumour region neurogenesis is incomplete as we do not observe fully formed ommatidia emphasizing the inhibitory effect of Dorsal on differentiation. In the prospective eye region, behind the morphogenetic frurrow, founder R8 neurons recruit further neurons through Receptor Tyrosine Kinase-MAPK signaling that potentially could add up with Ras^V12^-MAPK signaling to reach a higher signaling levels [49].

Extrinsic regulation of Toll signaling does have a role in the *Ras^V12^, scrib^-/-^ t*umor model for growth. The Toll-activating components implicated in tumour growth, PGRP-SA and ModSP, are part of a bacterial recognition alert system. The observed overexpression of PGRP-SA in *Ras^V12^*; *scrib^IR^* tumours may serve to locally increase the concentration of active cleaved Spz ligands, thereby stimulating Toll activation and promoting tumor growth. The source of activation governed by PGRP-Sa and ModSP remains unknown. As the posterior part of the disc experience higher levels of apoptosis, we considered whether apoptotic cell death and debris could active PGRP-SA/ModSP - driven Toll sigaling in this context through death-induced danger signals and bacterial component mimicry, but inhibition of death through DIAP1 expression did not alter NFkB levels.

Notably, previous studies have identified PGRP-SA, Cactus, and Dorsal as positive targets of the Toll pathway [34]. Furthermore, it has been shown that the CNS serves as a significant source of Spz ligands that regulates *Ras^V12^; lgl^-/-^* tumour migration tropism through Toll6 [33]. Hence, it is tempting to think that upon the onset of neurogenesis, tumour cells suddenly gain access to elevated levels of Spz concentration from the brain, significantly enhancing local Toll activation and subsequent PGRP-SA and Dorsal expression. This hypothesis may explain the spatial restriction of aberrant Dorsal expression to the posterior part of the tumor.

We have yet to assess whether DlA and DlB exert different functions within *Ras^V12^*; *scrib^IR^* tumours, as our knockdowns and overexpression transgenes targeted both isoforms simultaneously. Consistent with previous reports, DlB expression in the tumor context is restricted to cell types from the nervous system. Colocalization of DlB with pJNK in neurons that seem prone to die, allows us to speculate if both isoforms have distinct functions, that DlB may promote migration through JNK, while DlA may favor stemness and survival. Additionally, we cannot yet rule out the involvement of other NF-kBs, such as Dif and Relish in *Ras^V12^; scrib^IR^* tumours.

Using splicing variants may be a mechanism to diversify the range of NF-kB actions in various contexts in Drosophila, as it possesses only three NF-kBs (Dorsal, Dif, and Relish). In contrast, mammalian NF-kBs are not reported to have splicing variants, although they possess five different NF-kBs. To our knowledge, the concept of intratumoral NF-kB heterogeneity remains relatively unexplored in mammalian cancer biology and the fly model offer an experimentally accessible model to investigate this question *in vivo*. Future research efforts will be crucial in elucidating the importance and mechanisms underlying intratumoral NF-kB heterogeneity during malignant transformation.

The Ras^V12^, scrib tumor model engages five known inflammatory pathways during transformation. Ras^V12^ directly drives MEK-ERK and PI3K-AKT signaling while engaging Toll-NFkB, whereas loss of cell polarity through scribbled, initiates TNFR-JNK MAPK stress signaling and as a result, an Upd/IL6-JAK-STAT autocrine signaling loop altogether favoring malignant transformation.

In conclusion, the findings herein extends the understanding of how innate immune signaling is engaged by and cooperates with oncogenic Ras, and opens up for more detailed mechanistic studies in a genetically tractable preclincial tumor model with clear paralells to human cancer

## Material and Methods

### Drosophila lines

The EyMARCM system is known to generate flip-out clones in the adjacent optic lobes of the CNS in addition to the eye discs [50]. For assessing tumour cell migration from true carcinoma cells arising from the eye disc epithelium, we therefore used another genetic system designed in our lab, called EyaHost, that generates flip-out tumours exclusively in the EAD but not in the CNS (to be described in detail elsewhere).

Unless otherwise specified, crosses were conducted at 25°C by default; however, for enhanced knockdown efficiency, most knockdown experiments were carried out at 29°C, as detailed in the figures and legends.

- EyMARCM Drosophila lines:

Ey,flp; act>STOP>Gal4, UAS-GFP; FRT82B Tub-Gal80ts [32]

Ey,flp, UAS-Dcr2; act>STOP>Gal4, UAS-GFP; FRT82B Tub-Gal80ts

;Sp/CyO, Dfd-YFP; UAS-Ras^V12^ FRT82B scrib[2]^/^TM6C {Bilder, 2000 #23;Chen, 2012 #24;Perez, 2017 #22;Perez-Garijo, 2009 #25}

; UAS-PGRP-SA^IR^; UAS-Ras^V12^ FRT82B scrib[2]/TM6C (BDSC#60037)

; UAS-modSP^IR^; UAS-Ras^V12^ FRT82B scrib[2]/TM6C (VDRC GD#43972)

; UAS-pelle^IR^; UAS-Ras^V12^ FRT82B scrib[2]/TM6C (BDSC#50715)

; UAS-RFP^IR^; UAS-Ras^V12^ FRT82B scrib[2]/TM6C (BDSC#67852)

;UAS-Ras^V12^ ; FRT82B scrib[1]/TM6C [51]

;UAS-Ras^V12^ ; FRT82B

FRT82B scrib[1]/TM6C

FRT82B

- EyaHost Drosophila lines

; Eya-KD; QUAS-Gal4 (Eya-KD our own lab, QUAS-Gal4 BDSC#83132 {Baena-Lopez, 2018 #32})

QUAS-Ras^V12^ ; Eya-KD; QUAS-Gal4 (QUAS-Ras^V12^ from Peter Galant)

QUAS-Ras^V12^ ; Eya-KD, QUAS-scrib^IR^; QUAS-Gal4 (QUAS-scrib^IR^ from our own lab)

QUAS-Ras^V12^ ; Eya-KD, QUAS-scrib^IR^;

QUAS-Ras^V12^ ; Eya-KD, QUAS-scrib^IR^; QUAS-Gal4, UAS-Dcr2 (UAS-Dcr2 BDSC#24651)

Act>|STOP>|QF, QUAS-mCD8::GFP;; (Act>|STOP>|QF from our own lab, QUAS-mCD8::GFP BDSC#30001)

Act>|STOP>|QF, QUAS-mCD8::GFP; UAS-dorsal; (BDSC#9319)

Act>|STOP>|QF, QUAS-mCD8::GFP; UAS-modSP; (B.Lemaître {Dostalova, 2017 #33})

Act>|STOP>|QF, QUAS-mCD8::GFP; UAS-diap1^OE^; (BDSC#6657)

Act>|STOP>|QF, QUAS-mCD8::GFP;; UAS-cherry^IR^ (BDSC#35785)

Act>|STOP>|QF, QUAS-mCD8::GFP;; UAS-dorsal^IR-1^ (BDSC#32934)

Act>|STOP>|QF, QUAS-mCD8::GFP;; UAS-dorsal^IR-2^ (BDSC#27650)

Act>|STOP>|QF, QUAS-mCD8::GFP;; UAS-snail^IR^ (VDRC GD#6232)

Act>|STOP>|QF, QUAS-mCD8::GFP;; UAS-twist^IR-1^ (BDSC#51164)

Act>|STOP>|QF, QUAS-mCD8::GFP;; UAS-twist^IR-2^ (BDSC#25981)

Act>|STOP>|QF, QUAS-Tag2BFP;; genomic-dorsal::GFP (BDSC#42677)

Act>|STOP>|QF, QUAS-Tag2BFP;; genomic-cactus::GFP (VDRC fTRG 318145)

Act>|STOP>|QF, Ey-flp, QUAS-Tag2BFP; TRE-eGFP; (QUAS-Tag2BFP our own lab, TRE-eGFP BDSC#59010)

### Immunohistochemistry

Inverted larvae heads were fixed 30 min in 4% paraformaldehyde/PBS at room temperature on a shaker. After washes, fine dissection of single EADs (for early-stage tumours at Day 6) or whole cephalic complexes (at later stages) was conducted in PBS. The tissues were then incubated with a solution of primary antibody in 0,5% Triton X-100/PBS overnight at 4°C. Noteworthy, for primary antibodies that presented penetration defects (marked with * below), we incubated the tissues in 0,5% Triton X-100/PBS overnight at 4°C prior to primary antibody stainings. Similarly, secondary antibody stainings was done in 0,5% Triton X-100/PBS overnight at 4°C. Samples were then mounted in Vectashield or equivalent mounting media for imaging.

Primary antibodies were used at the following dilutions: anti-DlA mouse antibody (1:50, DSHB 7A4), anti-DlB rabbit antibody (* 1:1000, S.Wasserman [29]), anti-pH3 rat (1:200 abcam 10543), anti-Elav rat (1:50, DSHB 7E8A10), anti-cleaved Dcp1 rabbit (1:100, Cell Signaling 9578), anti-phosphoJNK mouse (* 1:100, Cell Signaling 9255 ) and anti-MMP1 mouse (* used in a mix 1:1:1, DSHB 3B8D12, DSHB 3A6B4 and DHSB 5H7B11). Secondary antibodies were all from Jackson ImmunoResearch.

### RNA sequencing and data analysis

Third instar larval eye discs of the indicated genotypes were manually isolated and RNA was extracted using Qiagen RNAmini kits. TruSeq RNA Library Prep Kit (Illumina) was used to prepare libraries, and sequencing was performed on an Illumina MiSeq system. Raw data were analyzed with the Tuxedo suite (RRID:SCR_013194) and reads were mapped to Drosophila genome release 5.2. Expression was recorded as FPKM: fragments per kilo-base per million reads.

### Imaging

Samples were imaged on Zeiss LSM880 and Zeiss LSM980 confocal microscopes as well as on Nikon HCI SoRa spinning disc microscope. They were further processed on FIJI.

### Tumour Volume Analysis

Z-stacks were acquired with a 3µm step for early-stage tumors (Day 6) and a 4µm step for late-stage tumors (after Day 6). Tumor volume was estimated using a FIJI Macro [52], which automatically performs threshold-based 3D reconstruction of the GFP+ signal, followed by volume measurements (3D Object counter plugin). Before 3D reconstruction, the FIJI Macro applies two consecutive Gaussian blur steps (sigma=2) to reduce noise.

### Image processing

- Fluorescence intensity measurements and heatmaps

Fluorescence intensity measurements were performed using a FIJI Macro, which subtracts background intensity to the signal of interest inside or outside the tumour, using a tumour mask generated from intensity thresholding. When comparing values across different conditions, they were normalized to the mean value of the control condition, ensuring the control mean is set at 1. In Fig 2B and Fig 2C, the tumours were split in two compartments “high” and “low”, according to arbitrarily set intensity thresholds (250 for DlA, 10000 for DlB). Normalized mean intensities were then automatically quantified within the different compartments: tumourHigh, tumourLow and TME. DlA and DlB intensity heatmaps were generated after background subtraction, using the Fire LUT in FIJI.

- Colocalization analysis

Colocalization analyses were performed using a FIJI Macro, which automatically computed the Pearson’s Correlation Coefficient (PCC) between two channels of a confocal image using the “coloc2” plugin. The PCC, ranging from -1 to 1, denotes the degree of colocalization between signals. A higher absolute value indicates stronger colocalization, with the sign indicating correlation direction (“-” for anti-correlation, “+” for correlation).

- Proliferation, differentiation and apoptosis ratio

The proliferation, differentiation and apoptosis ratio are defined as the area of the meaningful marker within the tumour (pH3, Elav and cDcp respectively) divided by the area of the tumour. These ratios were measured with a FIJI Macro which measures both the tumour area and the tumour marker area using an automatic-thresholding. For Fig 3A, 3B and 3C, measurements were conducted on single confocal pictures whereas, for S1 Fig, the apoptosis ratio was measured on the Z-projection of confocal Z-stacks.

- Migration score

Migration of the VNC and leg discs was manually scored on Z-projection (Max Intensity mode) of confocal Z-stacks.

### Statistical analyses

At least, two biological replicates were performed for each experiment. According to the normality and size of the datasets, appropriate statistical tests were applied using GraphPad Prism9 and are indicated in the figure legends. Graphs were also generated on GraphPad Prism9 and are presented as boxplots (10-90 percentile) or dot plots indicating the mean of the datasets. Statistical significance was categorized as follows: \**p* < 0.05, \*\**p* < 0.01, \*\*\**p* < 0.001, \*\*\*\**p* < 0.0001, and n.s when not significant.

## Acknowledgement

We thank Drs. Steven A. Wasserman and Amy Kieger, Bruno Lemaitre, the Bloomington Drosophila Stock Center; the Vienna Drosophila Resource Center and the Developmental Studies Hybridoma Bank for providing fly stocks and antibodies. We thank Alicia Alfonso Gomez for help with fly handling, experimental support and image acquisition. We also thank the Core Facilities for Advanced Light Microscopy and Advanced Electron Microscopy at Oslo University Hospital and the Microscopy Rennes Imaging facility of the UMS Biosit, a member of the national infrastructure France-BioImaging, supported by the French National Research Agency (ANR-10-INBS-04).

C.D, J.T.R., A.J., T.E.R and were financed by The Norwegian Research Council Toppforsk and Center of Excellence, CanCell grants #262652 and #276070

## Author contributions

CD and TER conceived the project and planned experiments. CD, AJ, JTR and H.J. performed experiments and analysis. H.J., and R.L. B. provided feedback to experimental approaches. T.E.R Is the leading principal investigator who together with C.D supervised research and edited the manuscript with input from all coauthors.

---

**S1 Fig.**
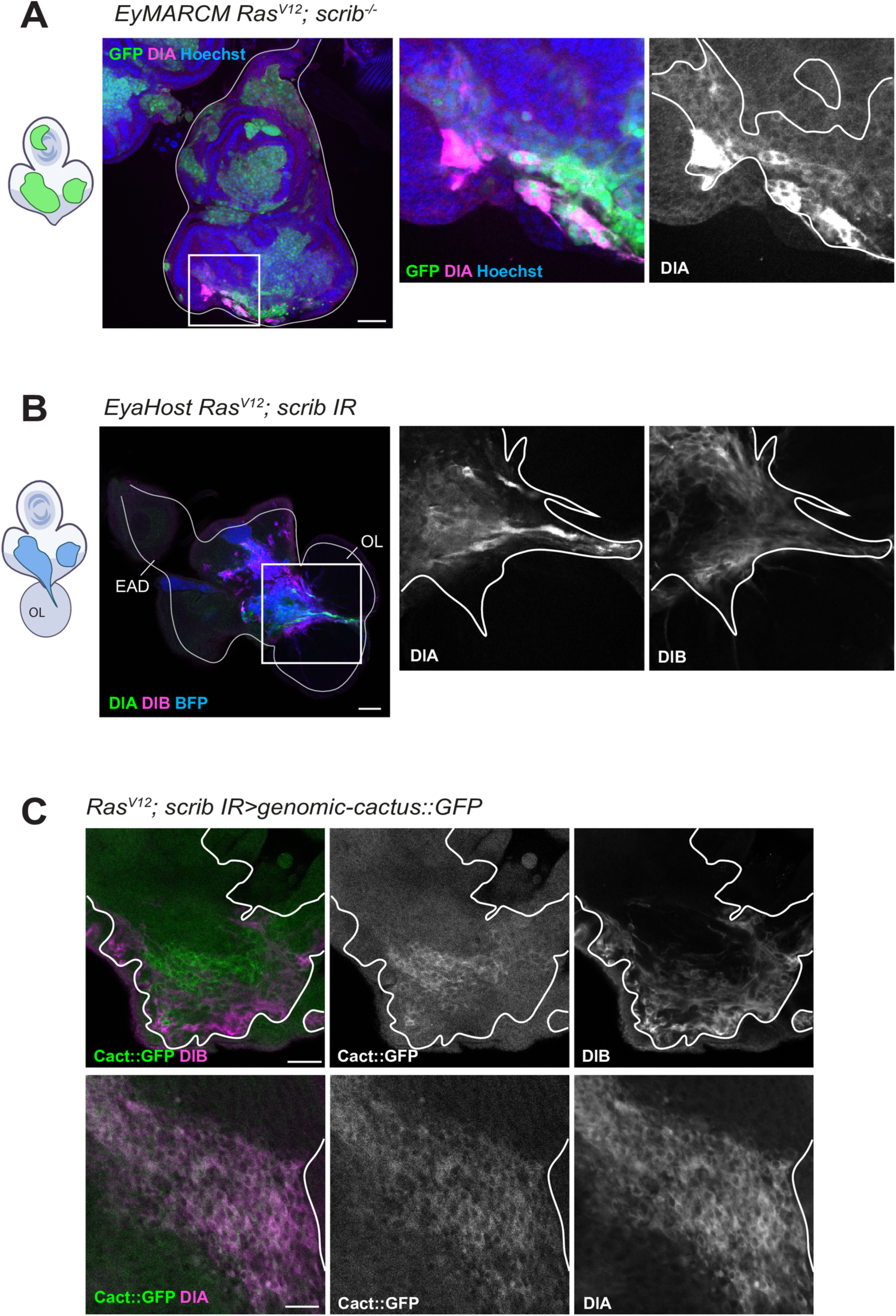
Localization of DlA, DlB and Cactus. (A) Representative confocal pictures of GFP-labelled *Ras^V12^; scrib^-/-^* tumours (n=10 EADs) at day 6, generated with the conventional EyMARCM genetic system and stained for DlA (magenta) and Hoechst (blue). Consistent with the data generated with our new EyaHost genetic system in Fig 2, DlA levels are aberrant in the tumours. The tumour is outlined with a straight white line. Scale bars=50µm. (B) Representative confocal pictures of GFP-labelled *Ras^V12^; scrib^IR^*tumours at day 6, generated with our novel EyaHost genetic system and stained for DlA (green), DlB (magenta) and Hoechst (blue). The tumour cells that already migrated inside the optic lobe (OL) of the CNS express high levels of DlA or DlB, suggesting their potential involvement in tumour cell migration. Scale bars=50µm. (C) Representative confocal pictures of BFP-labelled *genomic-cactus::GFP, Ras^V12^; scrib^IR^* tumours stained for DlB (magenta) at day 6 (upper line) (n=6 EADs) or stained for DlA (magenta) at day 8 (lower line) (n=6 EADs). The tumour is outlined with a straight white line. Scale bars=20µm.

**S2 Fig.**
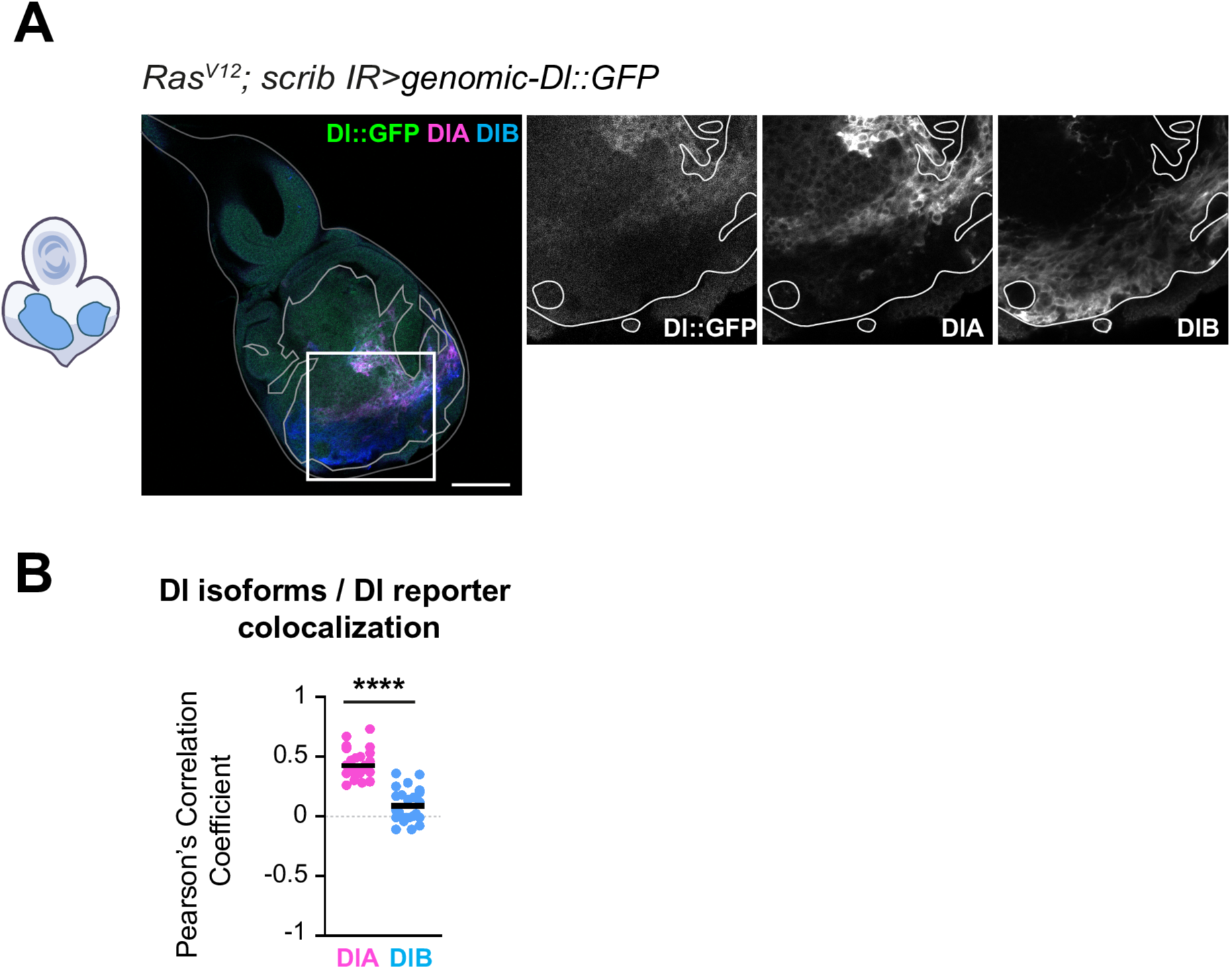
The endogenous Dl reporter used in this study is a DlA specific reporter. (A) Representative confocal pictures of BFP-labelled *genomic-dorsal::GFP Ras^V12^; scrib^IR^* tumours at day 6, stained for DlA (magenta) and DlB (blue). The tumour is outlined with the straight white line. Scale bars=50µm. (B) Colocalization analyses assessed with the Pearson’s correlation coefficient reveal a good colocalization between DlA and Dorsal::GFP (n=24, m=0,44, SD=±0,06) and no colocalization between DlB and Dorsal::GFP in the tumours (n=24, m=0,09, SD=±0,07) suggesting that this reporter is a DlA specific fusion protein. Statistical significance was determined with an unpaired T-test with Welch’s correction.

**S3 Fig.**
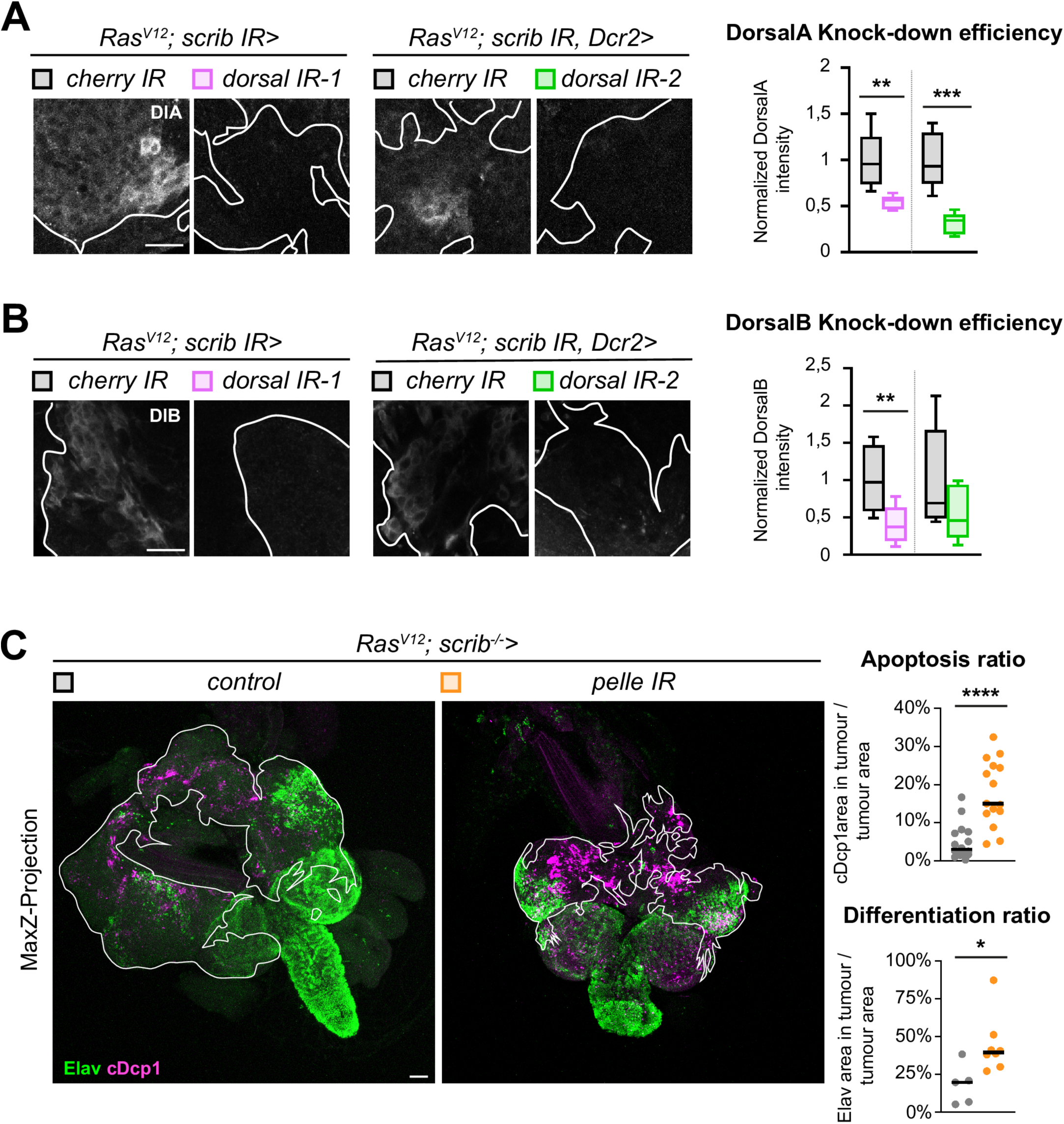
Dorsal knockdown efficiencies and pelle knockdown mimicks the function of Dorsal on differentiation and survival. (A) Representative confocal pictures and quantification of the mean DlA intensity of GFP-labelled *cherry^IR^ Ras^V12^; scrib^IR^* (n=6, m=1,00, SD=±0,14) and *dorsal^IR-1^ Ras^V12^; scrib^IR^* tumours (n=6, m=0,54, SD=±0,03) as well as *Cherry^IR^ Ras^V12^; scrib^IR^ Dcr2* control tumours (n=5, m=1,00, SD=±0,14) and *dorsal^IR-2^ Ras^V12^; scrib^IR^ Dcr2* tumours (n=6, m=0,32, SD=±0,05) at day 6 (29°C). Intensity normalization was done by subtracting the background intensity on each picture and finally dividing each data point by the mean intensity value for the control tumours for each replicate. Statistical significance was determined with a Mann-Whitney test. Scale bars=20µm. (B) Representative confocal pictures and quantification of the mean DlB intensity of GFP-labelled *cherry^IR^ Ras^V12^; scrib^IR^* (n=7, m=1,00, SD=±0.19) and *dorsal^IR-1^ Ras^V12^; scrib^IR^* tumours (n=9, m=0.40, SD=±0.12) as well as *Cherry^IR^ Ras^V12^; scrib^IR^ Dcr2* control tumours (n=5, m=1.00, SD=±0.31) and *dorsal^IR-2^ Ras^V12^; scrib^IR^ Dcr2* tumours (n=6, m=0.54, SD=±0.15) at day 6 (29°C). Intensity normalization was done by subtracting the background intensity on each picture and finally dividing each data point by the mean intensity value for the control tumours for each replicate. Statistical significance was determined with a Mann-Whitney test. Scale bars=20µm. (C) Z-projection of representative confocal pictures of GFP-labelled *RFP^IR^ Ras^V12^; scrib^IR^* control tumours and *pelle^IR^ Ras^V12^; scrib^IR^* tumours at day 6 (29°C), stained for Elav (green) and cDcp1 (magenta). The tumour is outlined with a straight white line. Quantification of the apoptosis ratio of *RFP^IR^ Ras^V12^; scrib^IR^* control tumours (n=16, m=4,97%, SD=±2,23%) and *pelle^IR^ Ras^V12^; scrib^IR^* tumours (n=15, m=17,84%, SD=±4,16%) at day 6 (29°C). Here, the apoptosis ratio is defined as the ratio between the cDcp1 area within the tumour and the area of the tumour itself. Statistical significance was determined with an unpaired T-test with Welch’s correction. Quantification of the differentiation ratio of *RFP^IR^ Ras^V12^; scrib^IR^*control tumours (n=5, m=18,04%, SD=±5,99%) and *pelle^IR^ Ras^V12^; scrib^IR^* tumours (n=8, m=44,14%, SD=±8,83%) at day 6 (29°C). Here, the differentiation ratio is defined as the ratio between the Elav area within the tumour and the area of the tumour itself. Statistical significance was determined with a Mann-Whitney test. Scale bars=50µm.

**S4 Fig.**
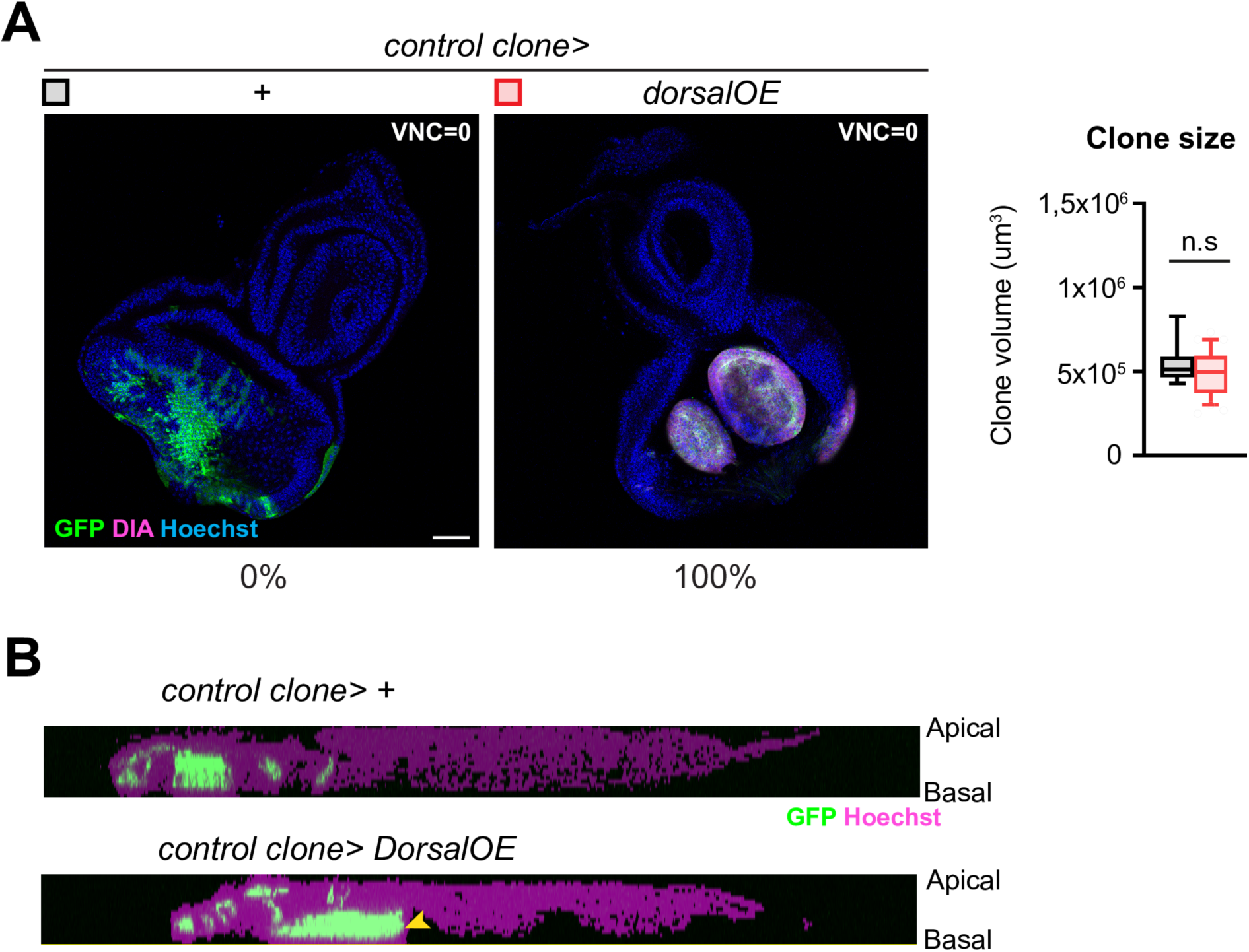
Dorsal is sufficient to induce dysplasia and delamination but not overgrowth and invasion. (A) Representative confocal pictures and quantification of the mean clone volume of GFP-labelled *wt* clones (n=20, m=5,59x10^5^µm^3^, SD=±0,69x10^5^µm^3^) and *dorsal^OE^* clones (n=30, m=4,87x10^5^µm^3^, SD=±0,66x10^5^µm^3^) at day 6, stained for DlA (magenta) and Hoechst (blue). Displasic clones are observed in 100% of the *dorsal^OE^*clones-bearing EADs. GFP-labelled cells are never found in the VNC as indicated with the VNC score values. Scale bars=50µm. (B) Reconstruction of an orthogonal view of *wt* clones and *dorsal^OE^* clones representing the EAD tissue (magenta) and the GFP-labelled clones (green). GFP-labelled cells extend from the apical to the basal side of the pseudostratified epithelium of the EAD of *wt* clones whereas some *Dorsal^OE^*clones spread basally (yellow arrow).

